# Neuropixels 2.0: A miniaturized high-density probe for stable, long-term brain recordings

**DOI:** 10.1101/2020.10.27.358291

**Authors:** Nicholas A. Steinmetz, Cagatay Aydin, Anna Lebedeva, Michael Okun, Marius Pachitariu, Marius Bauza, Maxime Beau, Jai Bhagat, Claudia Böhm, Martijn Broux, Susu Chen, Jennifer Colonell, Richard J. Gardner, Bill Karsh, Dimitar Kostadinov, Carolina Mora-Lopez, Junchol Park, Jan Putzeys, Britton Sauerbrei, Rik J. J. van Daal, Abraham Z. Vollan, Marleen Welkenhuysen, Zhiwen Ye, Joshua Dudman, Barundeb Dutta, Adam W. Hantman, Kenneth D. Harris, Albert K. Lee, Edvard I. Moser, John O’Keefe, Alfonso Renart, Karel Svoboda, Michael Häusser, Sebastian Haesler, Matteo Carandini, Timothy D. Harris

## Abstract

To study the dynamics of neural processing across timescales, we require the ability to follow the spiking of thousands of individually separable neurons over weeks and months, during unrestrained behavior. To address this need, we introduce the Neuropixels 2.0 probe together with novel analysis algorithms. The new probe has over 5,000 sites and is miniaturized such that two probes plus a headstage, recording 768 sites at once, weigh just over 1 g, suitable for implanting chronically in small mammals. Recordings with high quality signals persisting for at least two months were reliably obtained in two species and six different labs. Improved site density and arrangement combined with new data processing methods enable automatic post-hoc stabilization of data despite brain movements during behavior and across days, allowing recording from the same neurons in the mouse visual cortex for over 2 months. Additionally, an optional configuration allows for recording from multiple sites per available channel, with a penalty to signal-to-noise ratio. These probes and algorithms enable stable recordings from >10,000 sites during free behavior in small animals such as mice.

## Introduction

A major technical challenge for neuroscience is developing tools to record neuronal activity at large scale and across all relevant timescales (Chen et al., 2017; Seymour et al., 2017; Steinmetz et al., 2018; Hong and Lieber, 2019). Recent advances such as the Neuropixels probe leveraged CMOS fabrication methods to significantly expand the number and density of recording sites (Jun et al., 2017; Raducanu et al., 2017), allowing unprecedented recordings of large populations of neurons distributed across the brain at single spike resolution (Allen et al., 2019; Siegle et al., 2019; Steinmetz et al., 2019; Stringer et al., 2019b). The Neuropixels probe has seen rapid adoption and wide application in diverse species including mice (Evans et al., 2018; Vélez-Fort et al., 2018; Bennett et al., 2019; Kostadinov et al., 2019; Musall et al., 2019; Park et al., 2019; Schröder et al., 2019; Stringer et al., 2019a; Liu et al., 2020; Sauerbrei et al., 2020), rats (Krupic et al., 2018; Gardner et al., 2019; Böhm and Lee, 2020; Luo et al., 2020), ferrets (Gaucher et al., 2020), and non-human primates (Trautmann et al., 2019). Nevertheless, key barriers still prevent the recording of individual neurons stably over long timescales of weeks to months, of large-scale activity in small animals that are freely behaving, and of neurons packed densely in brain structures with diverse geometries.

The ultimate aim of chronic recordings is to record from the same neurons over days and weeks, but this goal has been difficult to achieve for large populations of neurons. Recording individual neurons stably over weeks or months is critical for studying the neural basis of processes that evolve over time, such as learning and memory, the dynamics of population coding, and neural plasticity. To provide prolonged high quality recordings, considerable effort has been devoted to the development of recording devices that are flexible (Fu et al., 2017; Chung et al., 2019; Musk, 2019) and/or <10 μm in size (Guitchounts et al., 2013; Luan et al., 2017; Egert et al., 2020; Welle et al., 2020) to minimize damage to tissue, but these approaches make insertion difficult and do not scale to large numbers of recording sites per inserted shank. Moreover, high quality signals can be recorded for more than eight weeks even with relatively rigid and larger devices such as wire tetrodes (Recce and O’Keefe, 1989; Dhawale et al., 2017), Utah arrays (Maynard et al., 1997; Chestek et al., 2011), and silicon probes (Okun et al., 2016; Jun et al., 2017; Muthmann et al., 2020; Schoonover et al., 2020). However, neither flexible nor rigid devices have been able to consistently record large numbers of identified individual neurons over weeks or months (Dhawale et al., 2017; Fu et al., 2017).

Rodents, especially mice, have become the dominant mammalian model for studying the neural basis of behavior, but their small size has made it challenging to record large populations of neurons during unrestricted movement. Implants that can be carried without impeding the behavior of a mouse must weigh less than ~3 g and span no more than 2 cm in height, and must connect with thin, flexible cables (or wirelessly). These limitations have precluded the use of many large-channel-count electrode arrays (Shobe et al., 2015; Rios et al., 2016; Scholvin et al., 2016). Even with the relatively small Neuropixels probe, which permits chronic recording in freely-moving mice, some impediments to movement have been observed (Juavinett et al., 2019), indicating the need for still smaller devices.

Finally, while single-shank silicon probes like Neuropixels achieve dense coverage along a line, some brain structures are more effectively recorded with other geometries. Several techniques including the Utah array, tetrode arrays, and microwire arrays offer the ability to sample across a plane approximately parallel to the brain surface (Recce and O’Keefe, 1989; Maynard et al., 1997; Saleh et al., 2019; Obaid et al., 2020; Sahasrabuddhe et al., 2020). However, to record in layered or deep structures such as isocortex, striatum, hippocampus, or superior colliculus, it can be ideal to densely sample a plane perpendicular to the brain surface (Shobe et al., 2015; Rios et al., 2016; Scholvin et al., 2016).

To address these challenges, we have developed the Neuropixels (NP) 2.0 probe. To record individual neurons stably across weeks and months, the probe has a denser, linearized geometry which allows for post-hoc computational stabilization of brain motion using a novel algorithm similar to image registration. To record in small, freely-moving animals, the probe and headstage were miniaturized to about one-third size, so that two probes and the one headstage needed to record them weigh ~1.1 g. This miniaturization was achieved without loss of channel count (384 channels / probe, where ‘channel’ refers to a signal processing and data transmission path). A 4-shank version of the probe can densely sample activity from a ~1 x 10 mm plane perpendicular to the brain surface. In total, 5,120 sites can be accessed across the four shanks, for 10,240 sites (768 of which are recordable simultaneously) using one headstage (where ‘site’ refers to a physical electrode location along a probe shank). New implantation hardware allows recovery and re-use of these implanted probes. Finally, we demonstrate a novel recording scheme in which two or more recording sites can be read out concurrently on a single channel, multiplying the number of sites simultaneously recorded. This strategy incurs a signal-to-noise ratio penalty but is suitable for recording spikes with high signal-to-noise ratio. This new device and our accompanying open source hardware and algorithms enable unprecedented stable, long-term, high temporal-resolution measurements of neuronal activity.

## Results

The NP 2.0 device design is miniaturized and optimized for long-term stable recordings in small mammals (Fig 1a). Like NP 1.0 probes, the NP 2.0 probe has a rigid base and 4 cm-long flexible cable that attaches to a headstage. The rigid area is shorter and narrower near the tip to facilitate close positioning of multiple probes. The headstage is also miniaturized, and serves two probes at once. Together, two probes plus a headstage weigh ~1.1 g, suitable for chronic implantation and freely-moving recordings in a mouse. The recording sites are denser with center-to-center spacing of 15 μm along the vertical dimension, compared to 20 μm for NP 1.0. Thus the number of sites per shank is 1,280 rather than 960, and there are both single- and 4-shank versions, with 5,120 recording sites in the latter. The recording sites are vertically aligned in two columns rather than staggered along the shank (Fig 1a), which is critical for the stabilization algorithm described below.

**Figure 1.**
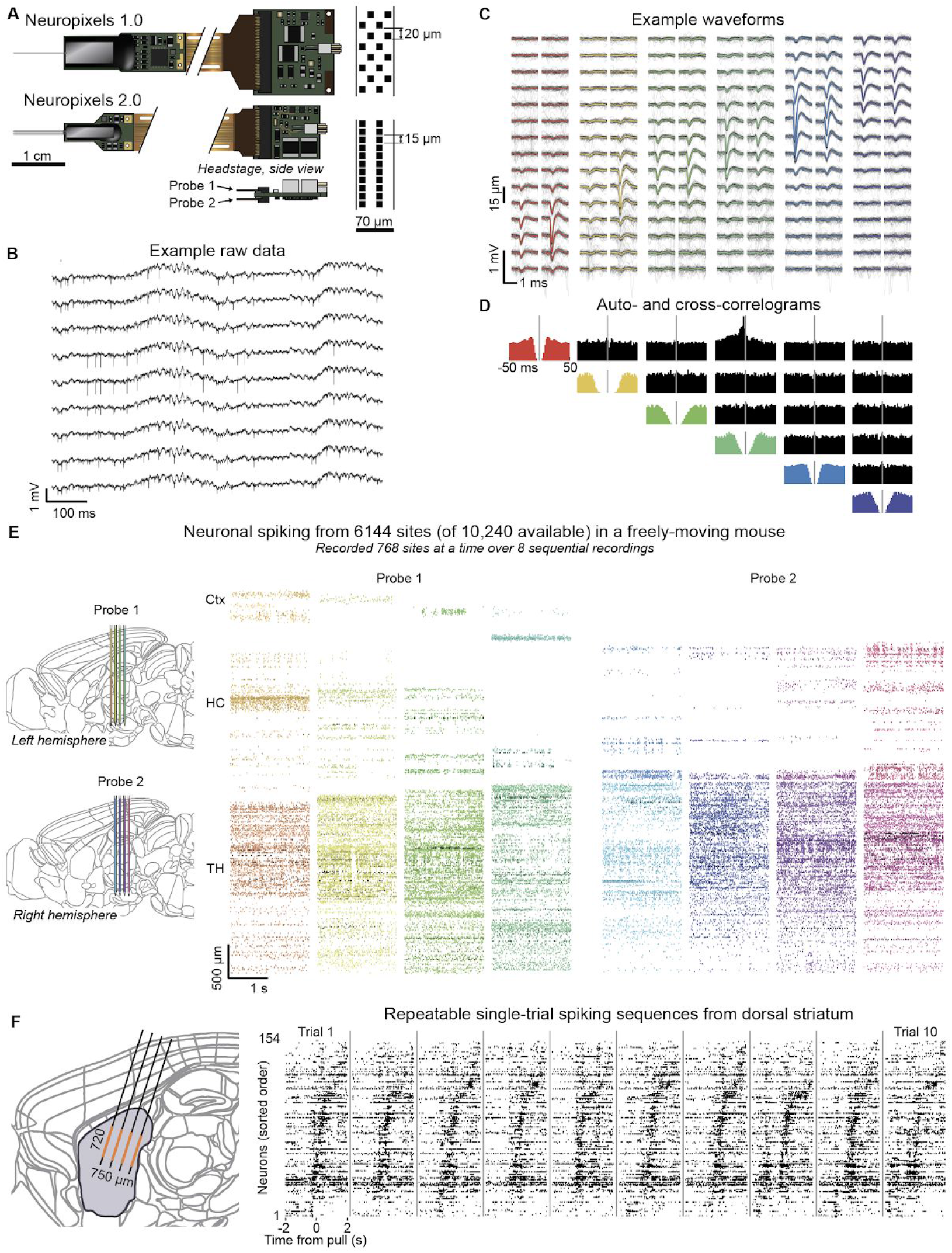
Neuropixels 2.0 devices are miniaturized and provide high-quality recordings across thousands of sites in vivo. (**A**) Comparison of the Neuropixels 1.0 (top) and 2.0 (bottom) devices. Neuropixels 2.0 have four shanks (or a single shank, not shown), miniaturized rigid base and headstage, and increased recording site density (right). They also allow for two probes to be attached to a single headstage (inset). (**B**) Example raw data traces show LFP and spiking signals recorded from 9 nearby channels within the olfactory bulb in an awake, head-fixed mouse. (**C**) Example spike waveforms from six selected neurons recorded on overlapping channels. The mean waveform (color) is overlaid on 50 randomly selected individual waveforms (grey). (**D**) Auto- and cross-correlograms (colored and black plots, respectively) of the example neurons from panel C, shown over a −50 to +50 ms window. (**E**) Example spiking rasters from two Neuropixels 2.0 probes chronically implanted in a single mouse, showing spikes recorded on 6,144 sites, out of the 10,240 sites available in total across the two probes. Each colored block represents spike times recorded from a ‘bank’ of 384 channels and plotted at the depth along the probe at which they occurred. Each probe could record one bank at a time, so that two banks (768 sites), were recorded simultaneously. The 6,144 sites were accessed by altering switches in software, and recording over 8 sequential recording epochs of 768 sites. (**F**) Dense local recordings from dorsal striatum in head-fixed mice performing a joystick-pulling task reveal reliable sequences of spiking activity on individual trials. Left, the 384 simultaneously recorded sites (orange) cover a plane 720 x 750 μm in extent, covering a significant proportion of dorsal striatum (purple). Recording location is illustrative, and does not represent a reconstruction from histology. Right, spiking raster from ten trials reveals characteristic spiking sequences across neurons, sorted for latency of peak response.

Despite being miniaturized, NP 2.0 probes each have 384 simultaneously recordable channels, with improved ADC resolution. The probes output a single wide-band 14-bit data stream (Fig 1b). The light sensitivity of NP 2.0 is similar to NP 1.0 (Supp Fig 1), and noise levels are slightly increased (from 5.4 to 7.2 μV root mean square voltage for the ‘alpha’ probes reported here, measured for the recording channel without electrode site noise, with further improvement expected in ‘beta’ version probes). Similar to recordings with NP 1.0, well-isolated individual neurons can be distinguished on overlapping channels (Fig 1c-d).

Hardware switches, programmable in software, allow rapid remapping of the recording channels to the recording sites, yielding recordings from thousands of sites per experiment (Fig 1e,f). The switches can be set again from software in less than one second without any physical manipulation of the probe or recording subject. In this way, thousands of sites can be recorded, 768 at a time, from a pair of probes with a single headstage in a freely-moving mouse (Fig 1e). By configuring the switches to record from a selection of sites across shanks, the 384 recorded channels of a single 4-shank probe can sample from sites densely covering a plane spanning 750 x 720 μm, an arrangement especially suited to structures of interest oriented in perpendicular to the brain surface (Fig 1f). This mode enables reliable observation of activity dynamics such as sequences in dense local populations.

NP 2.0 probes are optimized for high-yield chronic implants in small mammals. To confirm the quality of NP 2.0 recording characteristics over at least 8 weeks, as done for NP 1.0 probes (Jun et al., 2017; Juavinett et al., 2019; Luo et al., 2020), we implanted NP 2.0 probes in rats and mice in 6 labs. Out of 21 subjects implanted across the 6 labs, 20 implants were successful and recorded neurons until the experiment was ended at the discretion of the experimenter (see Table 1 for details). Data from an example recording shows maintenance of large-amplitude spiking activity over 8 weeks (Fig 2a) and consistent firing patterns across the depth of the probe for >44 weeks in one example recording (Fig 2b). Recording quality in most chronically implanted probes was maintained for at least 8 weeks across laboratories, as measured by the stability of total recorded firing rates (Fig 2c; for n=18/24 recordings the null hypothesis that firing rates were not declining with time could not be rejected; p<0.05, t-test for correlation coefficient over days, Fig 2d) and of spike sorted neuron count (Fig 2e; for n=19/24 recordings the null hypothesis that neuron count was not declining with time could not be rejected; p<0.05, t-test for correlation coefficient over days, Fig 2f). Seven of the 21 implants were performed with custom 3D printed fixtures (Supp Fig 2), which protected the electronics – i.e. the probe and headstage – throughout recordings and enabled probe recovery after the experiment. After recovery and cleaning, these probes were re-implanted in new subjects. In total, 7 out of 8 probes implanted with hardware suitable for recovery were recovered in working condition (Table 1). Finally, in one lab, recordings were made for more than 150 days in all implants (n=3), with a maximum time from implant to recording of 309 days, while retaining high firing rates, high neuron counts, and high quality individual neurons even at this late time point (Supp Fig 3). These results indicate that large populations of neurons can be reliably recorded with recoverable probes in both rats and mice, with little indication of an upper limit to the duration of these recordings.

**Figure 2.**
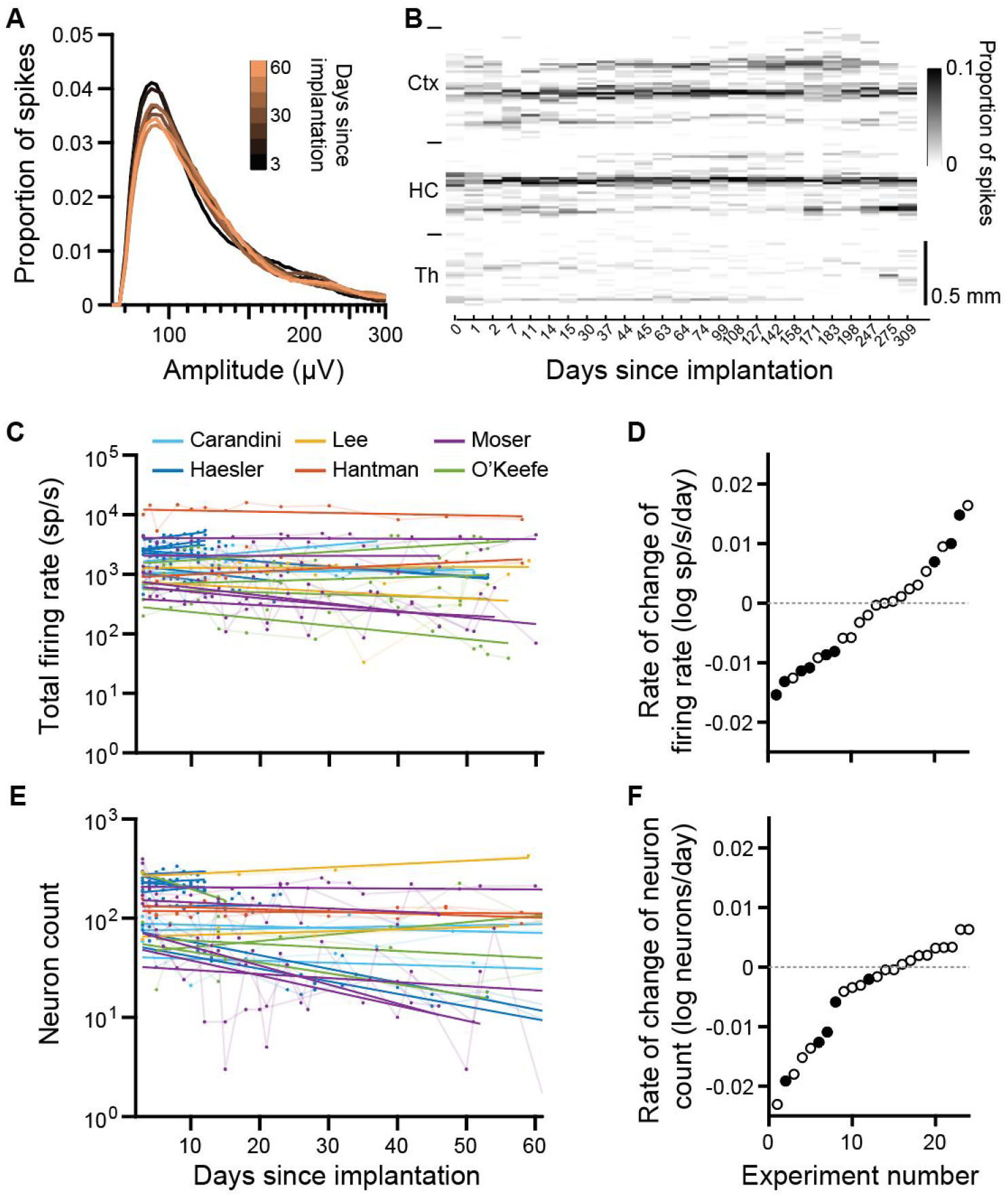
Chronic recordings with Neuropixels 2.0 probes maintained high yield for >8 weeks. (**A**) Stable distribution of spike amplitudes recorded on different days in the same subject. Spike amplitude distributions from each day are superimposed and color-coded by days since implantation. (**B**) Firing rates across channels are stable over nearly a year, in cortex (Ctx), hippocampus (HC), and thalamus (TH). Firing rates in spikes/s (sp/s) from each shank of the probe are separated by an orange line. Spikes are spatially binned across 15 μm. (**C**) Total firing rates over the course of 60 days for all probes used in this study. A linear regression line (in log_10_ units) was fitted to the total firing rate of each probe versus days since implantation. The color of each series represents data collected in different laboratories. (**D**) Rate of change in log total firing rate extracted from the linear fits (slope) of eac experiment in C. Each point represents one experiment. A rate of −0.01 log units per day indicates that over 100 days, the value declines by one log unit, i.e. a factor of 10. Filled dots represent significant correlations of the firing rate (or cluster count) with time. (**E**) Rate of change in log yield of spike-sorted neurons for each probe over the course of 60 days. (**F**) Same as D for neuron yields.

To record individual neurons stably over short and long timescales, it is necessary to maintain detection of spikes from the neuron over time and to match the spikes to the same unit. Classically, neurons are observed to decrease amplitude and disappear over individual recording sessions and over long timescales (days to months). We hypothesized that this amplitude decrease may be generally due to relative motion between the brain and the probe over time (Fig 3a, blue arrows). With a small number of recording sites a neuron that moves relative to the probe will be lost, but with a large number of recording sites the neuron will simply be recorded on different sites provided that brain-probe motion is in the direction along the length of the probe. We expect motion primarily along this axis because resistance to movement is much greater in axes perpendicular to the probe shank, along which the probe would have to sever the surrounding tissue to move. Indeed we commonly observed consistent shifts in spiking patterns along the length of the probe in many of our recordings (Fig 3b).

**Figure 3.**
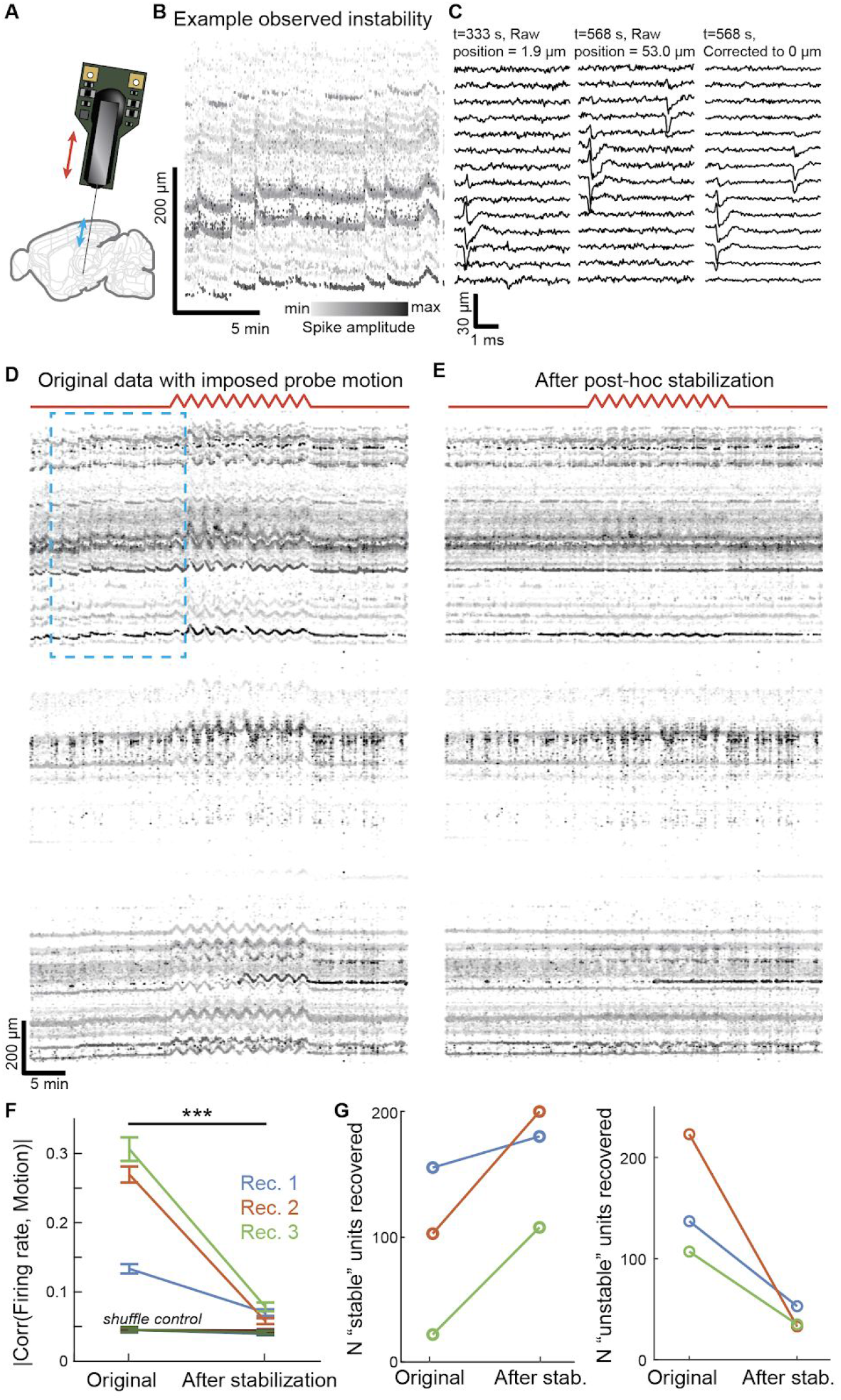
Post-hoc computational recording stabilization with an ‘image registration’ algorithm. (**A**) The brain moves relative to the probe primarily along the axis of the probe shank (blue arrows). In the test experiment, brain-probe relative motion was imposed by moving the probe up and down programmatically along the same axis with an electronic micromanipulator while recording (red arrows). (**B**) A spiking raster with spikes plotted at the position they occurred along the probe. Darker spikes have larger amplitude. Relative motion of neurons along the probe is observed across the whole probe over both relatively fast (<1 minute) and slow (~10 minutes) timescales, indicated by shared movement of the traces across depth. (**C**) Raw data snippets showing the stabilization approach. After motion was estimated, the raw data was re-sampled with spatial interpolation to shift the signals opposite to the direction of motion. The first sample shows a raw data segment from several channels where position was estimated near zero relative to the recording’s zero point. The second sample shows raw data from a later time point in the recording where position was estimated to be 53.0 μm from the zero point, or about three and a half channels shift. The third sample shows the results of correcting the raw data in the second sample to shift it to match position = 0. All spikes are shifted downward by this process, and the large spike to the left now aligns with the large spike from the first sample, presumably from the same neuron. (**D**) Spiking raster of a complete example recording with spikes plotted at the depth they occurred on the probe. The triangle-wave pattern of imposed probe-brain motion is shown in red, and is reflected in the movement of spikes along the probe during the middle of the recording. The blue dashed box shows the segment of data represented in panel B, which includes naturally occurring neuron motion. (**E**) Raster of spikes detected after applying the stabilization algorithm, showing stabilization of both imposed motion as well as naturally occurring motion. (**F**) The stabilization algorithm improved stability measured as the absolute value of the correlation coefficient between firing rates and probe-brain motion. (**G**) The stabilization algorithm improved yield of neurons whose firing rates had no correlation with the imposed motion (“stable”) and reduced the number whose firing rates correlated with the motion (“unstable”).

We therefore asked whether spikes from individual neurons were preserved and detectable even during brain-probe relative motion, and if so, whether we could correct for the effects of this motion. To test this, we devised an approach to give us ground truth knowledge of the motion of the probe relative to the brain, in order to assess whether the effects of the motion could be computationally stabilized. We performed acute recordings in awake, head-fixed mice during which the probe was moved up and down programmatically using an electronic micromanipulator to impose a fixed and known pattern of brain-probe relative motion on the recording (Fig 3a, red arrows; 10 cycles of triangle wave movement with amplitude 50 μm and period 100 s / cycle).

We then devised a novel unsupervised algorithm to stabilize recordings post-hoc, and we tested it on these ground-truth datasets with known movement. The algorithm determines the motion over time from the spiking data and stabilizes it with spatial resampling of the original raw data, as in image registration (Fig 3c; see Methods). Before applying the post-hoc stabilization algorithm, datasets with imposed motion clearly reflected the triangle wave pattern (Fig 3d). Applying the algorithm to the raw data and then re-detecting spikes demonstrates that the algorithm removed the relative motion between the brain and the probe, resulting in stable patterns of spiking activity over time (Fig 3e). We assessed whether each neuron’s spikes were detected throughout the motion by computing a correlation coefficient between each neuron’s firing rate and the probe position. If the spikes from a neuron are lost when the neuron is located at a certain position along the length of the probe, then the observed firing rate would decrease in a way that is correlated with the time course of the imposed motion. The post-hoc stabilization algorithm significantly reduced the absolute value of correlation between firing rate and probe position, decreasing it to near chance levels (Fig 3f; 0.224±.007 mean±s.e.m. before correction, 0.067±.003 after stabilization, chance level .042±.001; two-way ANOVA, main effect of stabilization p<10^-10^). Moreover, the algorithm markedly improved yield of neurons with stable firing rates (n=156, 103, and 22 stable units without stabilization versus n=181, 201, and 108 stable units respectively with stabilization; corresponding but opposite changes in the number of unstable units; Fig 3g). Applying post-hoc stabilization also improved firing rate motion correlation and neuronal yield in data obtained under the same paradigm with NP1.0 probes, though the firing rate motion correlation after stabilization was significantly higher (i.e. worse) than that achieved in NP2.0 datasets (0.097±.005 mean±s.e.m., two-way ANOVA, main effect of probe type p<10^-10^; data not shown). Stabilization was presumably more successful in NP2.0 probes because of the vertically aligned sites and smaller gaps between sites (Fig 1a), which together increase the spatial resolution of sampling along the direction of motion.

Relative movements between the brain and probe occur not only on a fast time scale but also across days. Given that NP 2.0 probes record from neurons stably even as the brain moves relative to the probe, we reasoned that we could apply a similar approach to data recorded across multiple sessions over weeks or months. As in the acute situation, we observed that the spiking activity appeared to represent the same patterns but shifted in depth across weeks (Fig 4a).

**Figure 4.**
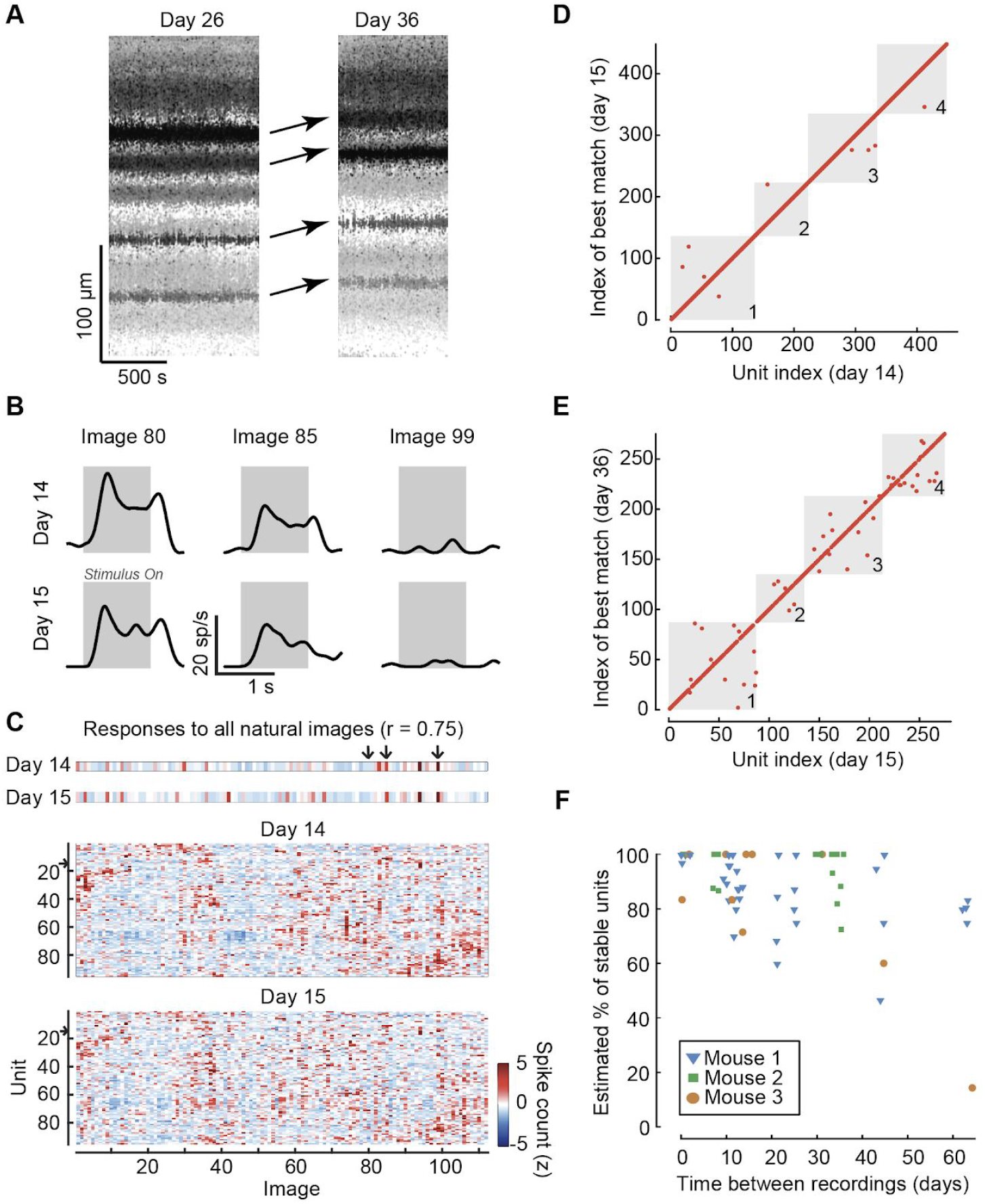
Successful tracking of neurons across days and weeks. (**A**) Data from an example mouse with a chronic NP 2.0 probe in visual cortex, showing significant drift between recordings on consecutive days. Plotting conventions as in Fig 3b. (**B**) Peri-stimulus time histograms (PSTHs) of an example neuron for three images (n=5 repetitions each) eliciting different levels of response. The visual stimulus was presented for the time period marked by the grey box. (**C**) Top: Average spike count response of the same unit to all the natural images presented to the animal (the three example images from b are indicated by arrows) on two consecutive days (z-scored). The two responses have a correlation of 0.75. Bottom: Example average spike count response of all units with visual fingerprint on one of the shanks on two consecutive recording days, with both neurons and images in sorted order according to similarity of responses. Color bar: z-scored response. (**D**) In order to gauge unit stability by assessing visual response stability, each unit’s visual fingerprint on the first day was matched with its own fingerprint and with the fingerprint of the closest other unit (not necessarily labeled with a consecutive index) on the second day. The majority of units matched to themselves (points on the diagonal, 439/448), but in a few cases units matched better with the visual fingerprint of their neighbor (red points off the diagonal, 9/448). Each grey square delineates the units belonging to one of the four shanks, indicated by the numbers. (**E**) Same format as D, for two recordings that are three weeks apart (229/275 are matches). (**F**) Summary of stability of well-isolated units across 26 spliced pairs of recordings in three mice. Each point represents a single shank, and data from each subject is shown by a different symbol. In total, 1748 well-isolated units with visual fingerprints were analyzed. The estimated percentage of stable units is calculated as 2Prtmateh) – 1 where Pr(match) is the probability that a unit’s visual fingerprint matched more closely than the nearest neighbor on the two days (see Methods for derivation). For presentation purposes only, points were jittered along the x-axis. Note that the interval between implantation and the first recording in each pair of recordings was variable, in some cases exceeding 6 months.

To track neurons across sessions, we implemented a version of the motion stabilization algorithm described above. Unlike the dynamic stabilization, the version used here inferred and corrected the shift only at the point where datasets from the two days are spliced together, i.e. at the end of the first and start of the second dataset (see Methods). Following this step, spikes were sorted together across the splicing point, using only the shapes and spatial footprints of the waveforms. Thus, spikes were joined into single clusters across days without reference to their functional properties.

With this improved algorithm for motion stabilization, we were able to record from the same neurons in visual cortex across days. To establish a ground-truth metric for whether a group of spikes (a “unit”) recorded across two different days corresponded to a single neuron, we relied on the fact that neurons in rodent primary visual cortex have visual responses unique amongst their neighbors (Ohki et al., 2005). Here, we used a battery of 112 natural images to establish a visual fingerprint of a large subset of units (Figure 4b,c). We then assessed whether the algorithmically-tracked units represented the same neuron by determining whether their responses to our battery of images across two sessions were more similar to each other than to the responses of the nearest other unit. This procedure revealed that most units are successfully tracked across days (Fig 4d) and weeks (Fig 4e). For recording sessions that are 16 days apart or less, an estimated 93%±9% of well-isolated units were successfully tracked in time (mean±st. dev. across shanks; n=1110 units, 36 shanks, 15 recordings in 3 subjects; see Methods; Fig 4f). Across sessions separated by 3-9 weeks, we could still successfully track 83%±20% of the well-isolated units (n=638 units, 30 shanks, 11 recordings in 3 subjects, Fig 4f). In one of the three subjects we observed a case of a loss of almost all of the tracked units, which we speculate may have been due to a non-coaxial shift of the probe relative to the brain. We did not attempt to track units across this discontinuity event. However, even in this mouse we were able to track units if both recording days were on the same side of the discontinuity. Recording with NP2.0 and applying our stabilization algorithm therefore permits the study of neuronal and population coding over timescales of learning and plasticity.

Finally, we reasoned that the high signal-to-noise ratios (SNRs) of many recorded neurons may allow a novel strategy for increasing recording coverage, in which the signals from multiple distant recording sites are combined on a single recording channel (Fig 5a). From physical considerations, connecting two distant sites to one electrical line and one recording channel ought to average the signals at the two sites, in principle allowing for recording from twice as many (or more) sites as there are channels on the device. This strategy would result in a diminution of signal magnitude by a factor of 2 and changes the noise level according to the equation:

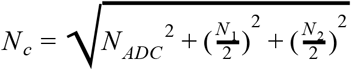

where *N_c_* is the noise level when recording on banks 1 and 2 combined, *N_ADC_* is channel noise from the recording system, and *N*_1_ and *N*_2_ represent noise from biological and physical sources at each electrode site (assumed uncorrelated). The SNR would thus decrease by a factor of 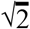 (in the limit where *N*_1_ and *N*_2_ substantially exceed *N_ADC_*). However, although the SNR of each neuron will decrease, the number of recorded sites will double, yielding a viable strategy for scaling recording beyond limits on the number of recording channels.

**Figure 5.**
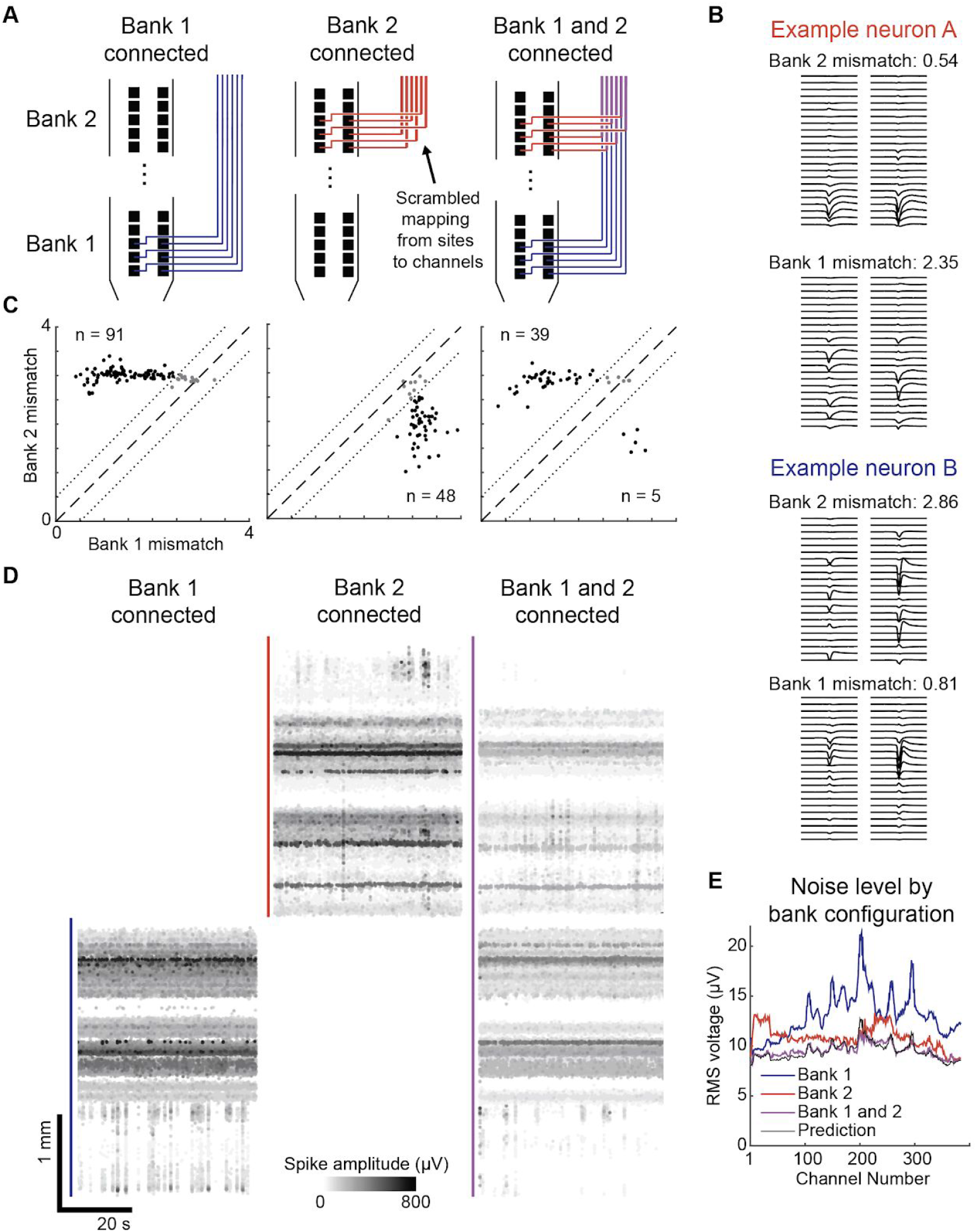
Recording from twice as many sites as the number of recording channels. (**A**) Sites from multiple banks connect to a single set of recording channels, and switches control which sites connect to which channels. 10 sites out of 384 are illustrated for each bank. Right, when both sets of switches are closed, the recorded signal will combine the signals at each bank of sites (indicated by purple color). In bank 2, the mapping from sites to channels is scrambled relative to bank 1. (**B**) Due to the scrambled mapping, waveforms of recorded neurons appear correctly clustered only when interpreted as arising from one of the two banks. Each panel shows the waveforms of the example neuron arranged spatially according to either the bank 2 (top) or bank 1 (bottom) channel mapping. (**C**) The mismatch score for banks 1 and 2 under conditions when bank 1 was connected alone (left), bank 2 was connected alone (center), or both banks were connected (right). The dotted lines demarcate the region where mismatch scores differed by less than 0.5 between the banks; units in this range (grey points) were excluded as being uncertain. (**D**) Spiking raster (conventions as in Fig3b) from recordings with all three configurations. In the condition with bank 1 and bank 2 recorded together, spikes are plotted at their inferred locations based on the mismatch score of their source template. Spike amplitudes are lower by a factor of two with both banks connected. (**E**) The noise level on each site is plotted under each recording configuration. The prediction of the combined bank 1 and 2 noise level using the equation given in the text is shown in black.

We designed the single-shank version of NP 2.0 to allow for connecting multiple sites to a single recording channel, with a modification to the channel mapping that allows for reconstruction of the locations of the original recorded neurons. In particular, we designed the mapping between channels and sites to be scrambled from one bank of 384 sites to the next (Fig 5a). Since nearly all recorded neurons are visible on more than one recording site, the pattern of observed waveforms across channels from one neuron will form a spatially compact group only when interpreted as arising from one bank, and not the other (Fig 5b). To classify recorded neurons to channel banks, we devised a “mismatch score” to describe how dispersed the waveforms are across sites under each bank’s channel mapping, with low scores indicating compact waveforms. This procedure could reliably identify which bank each neuron was recorded on, in that recordings made truly on just one bank resulted in nearly all neurons correctly classified as arising from the recorded bank rather than from the other (Fig 5c; 393/394, 99.8% of neurons, n=3 recordings; see Methods for detailed criteria).

Using this strategy, we could reconstruct the pattern of spiking activity across 768 recording sites in a single recording session despite recording with only 384 channels (Fig 5d). The spikes recorded in the double-bank configuration were indeed lower amplitude than those in the single bank configuration by a factor of 2. Moreover, the noise level on the combined channels closely matched the prediction of the equation given above (r = 0.92 and 0.95 in two recordings; Fig 5e). In our recordings, the empirical SNR of spikes recorded in the double-bank configuration was 63.5 ± 0.3% that of the single bank configuration (mean±s.e.m., n=1536 recording sites, 2 recordings). As a result, the yield of sortable single neurons was less than the total yields when recording each bank separately (summed yields of 215, 139, and 40 neurons for separate banks versus 75, 44, and 20 for combined banks). Nevertheless, these neurons were recorded across a span of the brain twice as long as that covered by a single bank of sites. This is the first demonstration of truly simultaneous recording across more sites than available channels, a recording strategy that is suitable for capturing neurons with large SNR over a large spatial extent.

## Discussion

In summary, we have demonstrated a suite of novel electrophysiological tools: a miniaturized high-density probe; recoverable chronic implant fixtures; software algorithms for fully-automatic post-hoc computational stabilization; and a strategy for extending the number of recorded sites beyond the number of available channels. We have presented experiments that validate the quality of the recordings, the stability of the recordings over timescales of months, and the reliability of their use chronically in six different laboratories. Finally, we provide ground-truth proof of the efficacy of stabilization on both short and long timescales. Together these tools enable an order of magnitude increase in the number of sites that can be recorded over long timescales in small animals such as mice, and the ability to record from them stably.

While our approaches for computationally stabilizing recordings achieved marked improvements over prior state of the art, as well as improvements relative to processing the same datasets with previously available algorithms (Figs. 3, 4), we nevertheless did not completely eliminate indications of instability (Fig 3f) nor could we record every neuron stably over long timescales (Fig 4d). The remaining instability could be due to underlying experimental factors: mechanical forces in experiments with imposed probe motion could alter neuronal firing of nearby neurons; and, neuronal death or true changes in visual response properties (Deitch et al., 2020) could result in failure to track neurons over weeks. Another possibility is that the remaining instability could be due to imperfect spatial sampling of the NP 2.0 probe. In biophysical models of neurons’ extracellular potentials (Gold et al., 2006), some features of these potentials are predicted to be smaller than the spatial sampling density of this probe (15 μm), raising the possibility that applying the algorithms developed here to data from probes of still higher density (Fiáth et al., 2019) may in the future yield even better solutions to these challenges.

## Supplemental Figures and Table

**Supplementary Figure 1.**
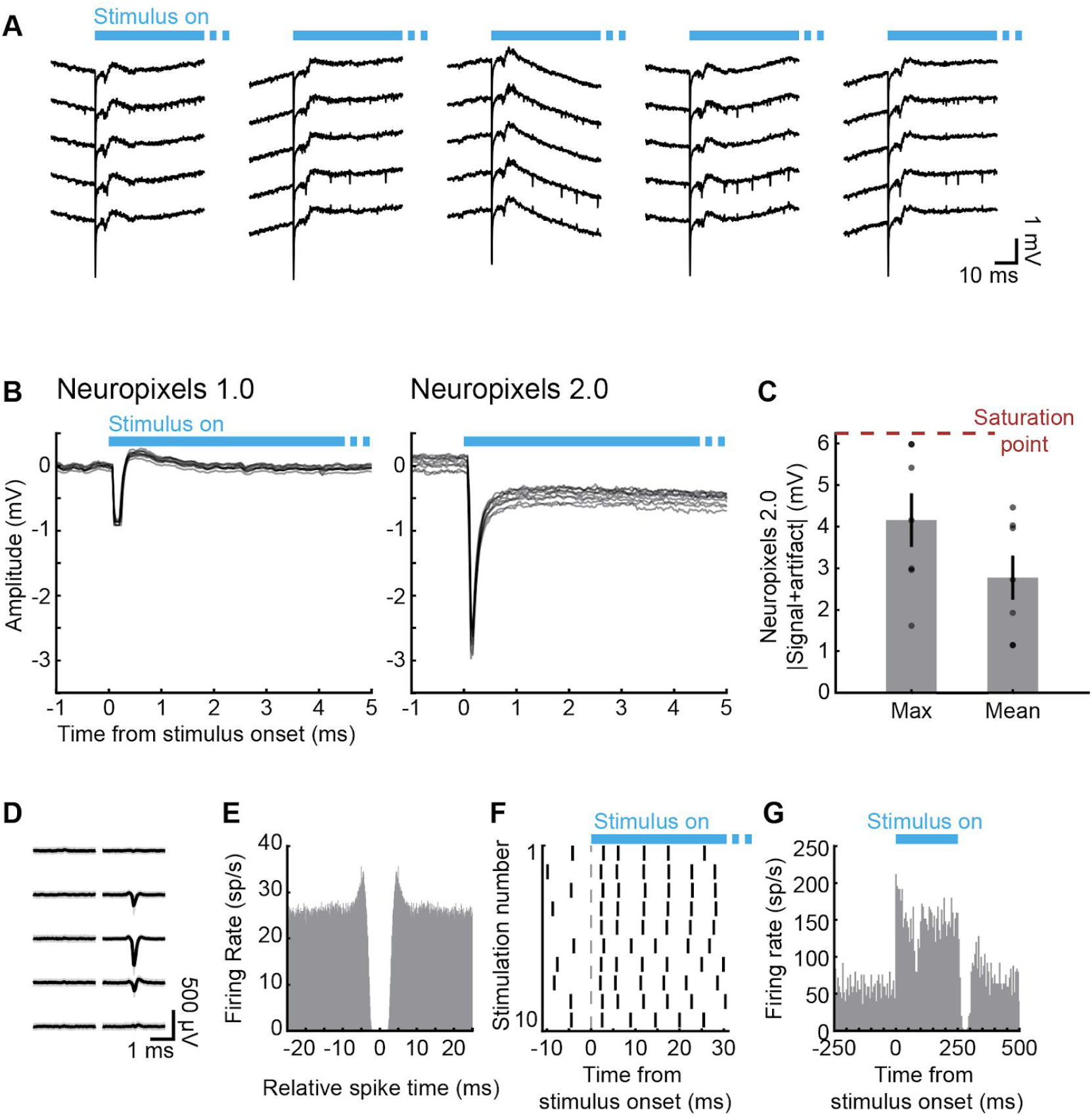
Optogenetic activation of neurons recorded with NP2.0. (**A**) Raw data traces show LFP and spiking components recorded from 5 nearby channels in the cerebellar cortex in an awake, head-fixed mouse. Optogenetic stimulation fiber was placed on the brain surface and channels shown are approximately 750 μm below the brain surface. (**B**) Optogenetic stimulus artifacts recorded with NP1.0 probe (using gain=500, left) and NP2.0 probe (right) showing that comparable levels of stimulation saturate NP1.0 probes used at recommended gain settings but not NP2.0 probes. Signals from 10 adjacent channels recorded on a single trial are overlaid. (**C**) Maximal (left) and mean (right) optogenetic stimulus-evoked signal amplitude recorded with NP2.0 probes (n = 7 recordings, 7 mice). Optogenetic stimulation never saturated NP2.0 probes. (**D**) Example waveform of a neuron responsive to optogenetic stimulation in an L7-cre; Ai32 mouse (i.e., a Purkinje cell). (**E**) Autocorrelogram of the neuron shown in panel D measured from spikes recorded during an initial baseline period. (**F**) Spike raster of the neuron shown in panel D in response to optogenetic stimulation (after synaptic cocktail application). Note that the neuron responded reliably within ~3 ms of stimulus onset. (**G**) Peri-stimulus time histogram of the neuron shown in panel D showing full response to optogenetic stimulus.

**Supplementary Figure 2.**
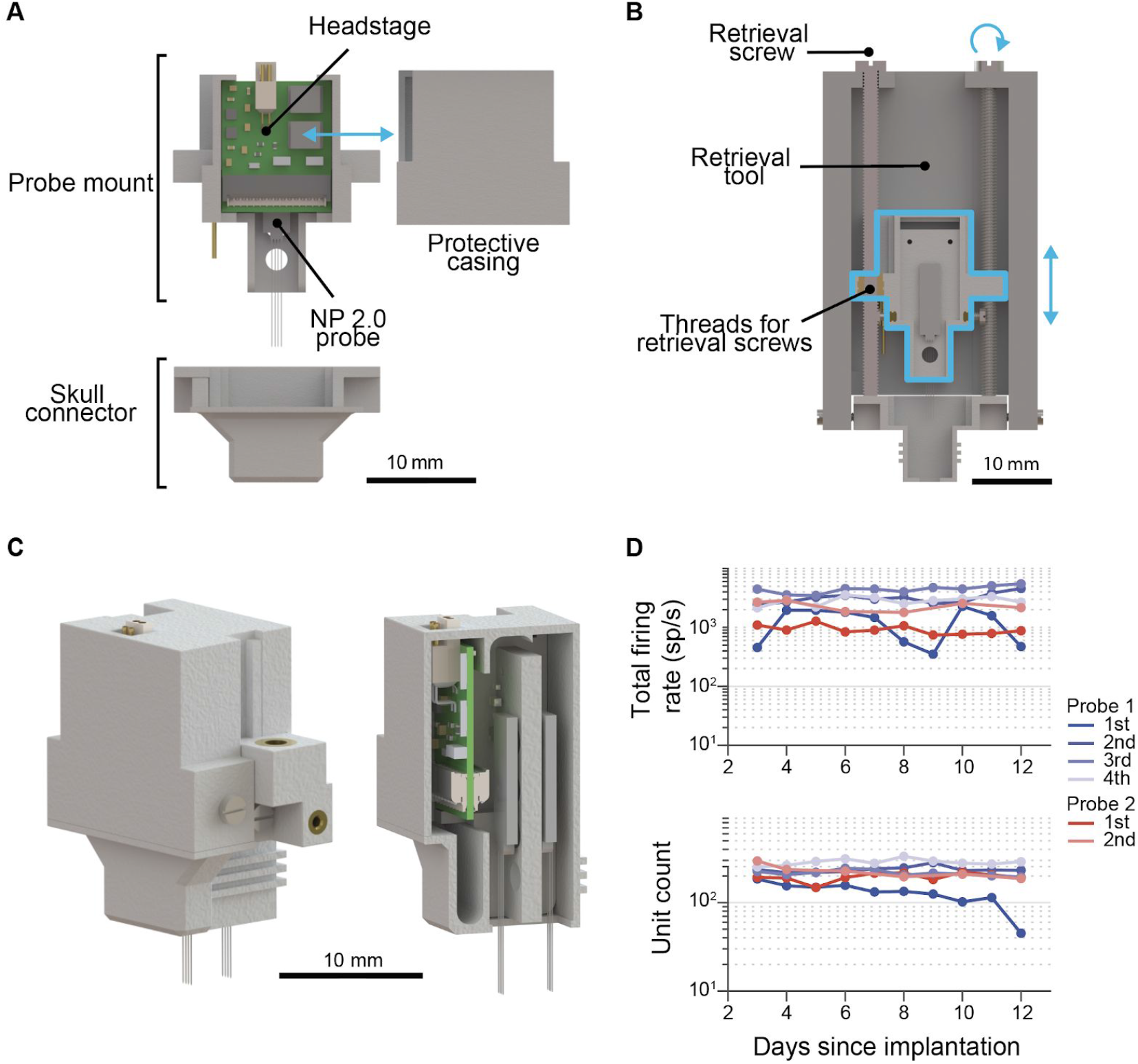
A custom implantation fixture enables reusing probes in chronic experiments. (**A**) Schematic overview of the single-probe fixture. (**B**) Retrieval mechanism. This tool is used to help extract the probe after the conclusion of the experiment, and is not carried by the mouse during the experiment. (**C**) Rendering and cross section of dual-probe fixture. Total weight of this fixture is 2.76 g. (**D**) Consistent high unit yield with reimplanted probes. Two probes (probe 1, blue, probe 2, red) were separately implanted into olfactory cortex 4 and 2 times, respectively, in different mice. Each implantation resulted in similar total firing rates (top) and unit yield (bottom), obtained over the 12 day period of the experiment. These results demonstrate that the fixture enables consistent, stable recordings with reused probes.

**Supplementary Figure 3.**
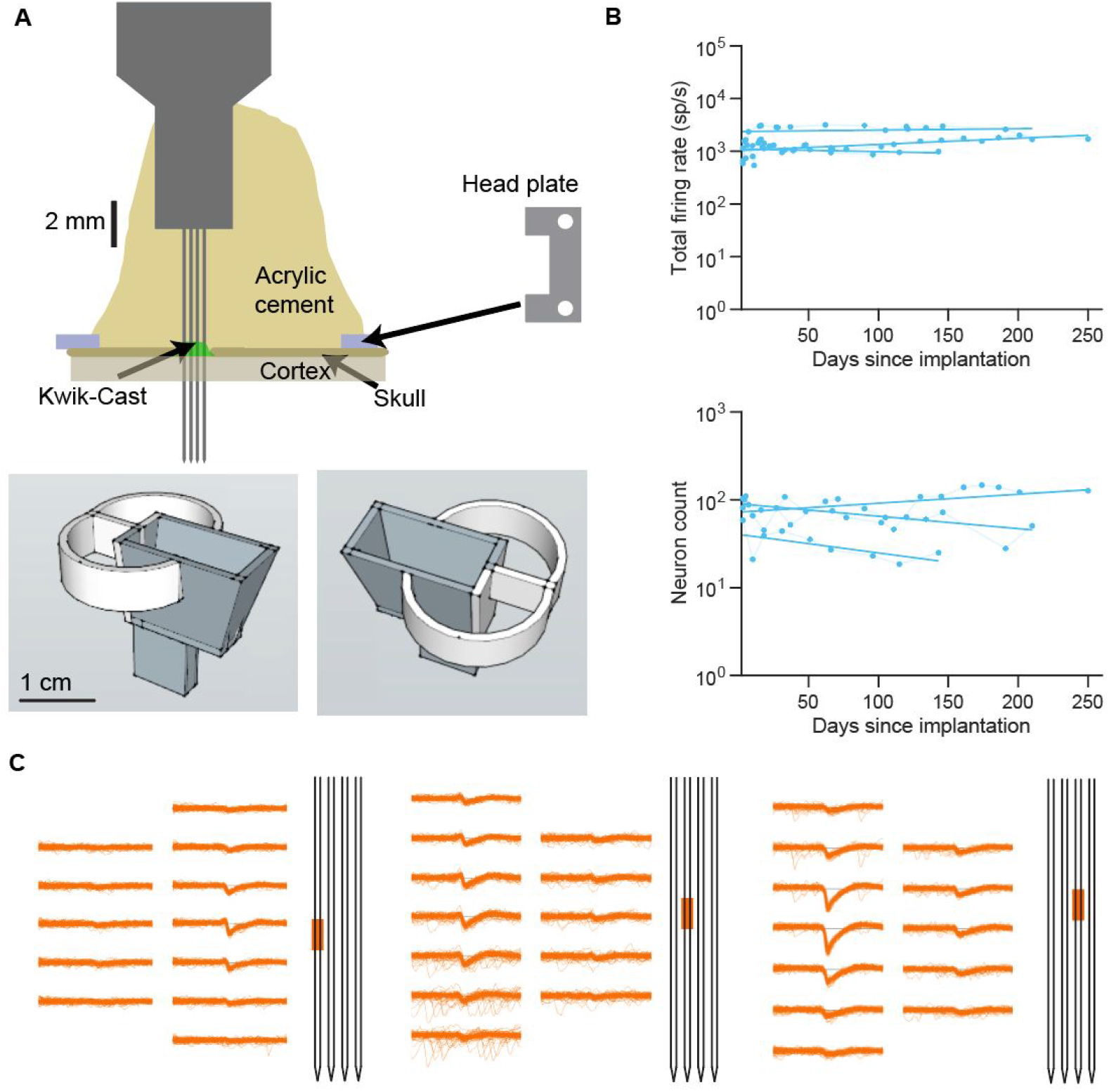
Chronic recordings with NP2.0 for >200 days. (**A**) Top: Schematic of the implantation (for a 4-shank NP 2.0 probe). The shanks of the probe were inserted approximately 5 mm into the brain, through primary visual cortex. The brain was subsequently protected by a thin layer of Kwik-Cast. The probe was then cemented to the skull. The head-plate on the right is not shown to scale. Bottom: Schematic of a custom 3D printed holder. The holder was lowered after the probe was securely cemented to the skull, so that the probe PCB was protected by the semi-circular rim (shown in white), then the central part of the holder (shown in grey) was cemented to the skull above the frontal cortex. (**B**) Total firing rates and unit yields over the course of 150-250 days for the 3 animals with primary visual cortex implants (performed as illustrated in A). A linear regression line (in log space) was fitted to the change in total firing rate of each probe and unit yield versus days since implantation. (**C**) Example waveforms of neurons in primary visual cortex from three shanks of the probe, recorded 237 days after implantation. Orange marks on the cartoon of the 4-shank probe depict the region where the sample channels were located.

**Supplementary Figure 4.**
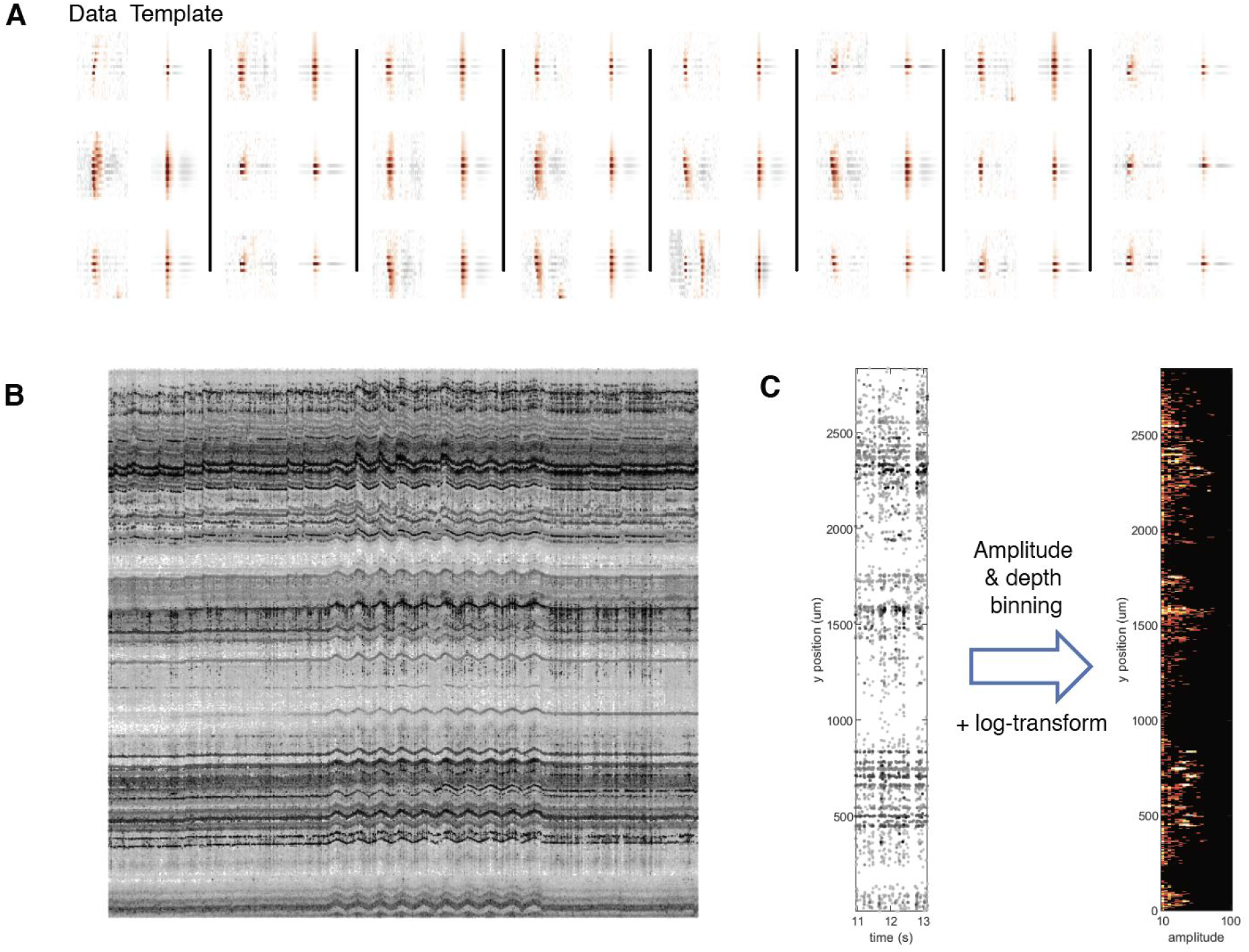
Spike detection for the motion stabilization algorithm. (**A**) Randomly-chosen examples of 24 spikes detected from the recording shown in the main Figure 3, together with the generic templates that were used to match these spikes in the raw data. Red/gray hues indicate negative/positive potentials. The nearest 21 channels to each channel peak are shown and 61 time samples. (**B**) Example drift map for the entire ~45 min recording, including the period of manipulator movement from 1000 s to 2000 s. Same conventions as main Figure 3 (darker points indicate larger amplitudes). (**C**) (left) Spikes from a single 2 s batch of data, i.e. zoom in of panel B. (right) Conversion of the spikes in this batch to an amplitude/depth histogram.

**Supplementary Figure 5.**
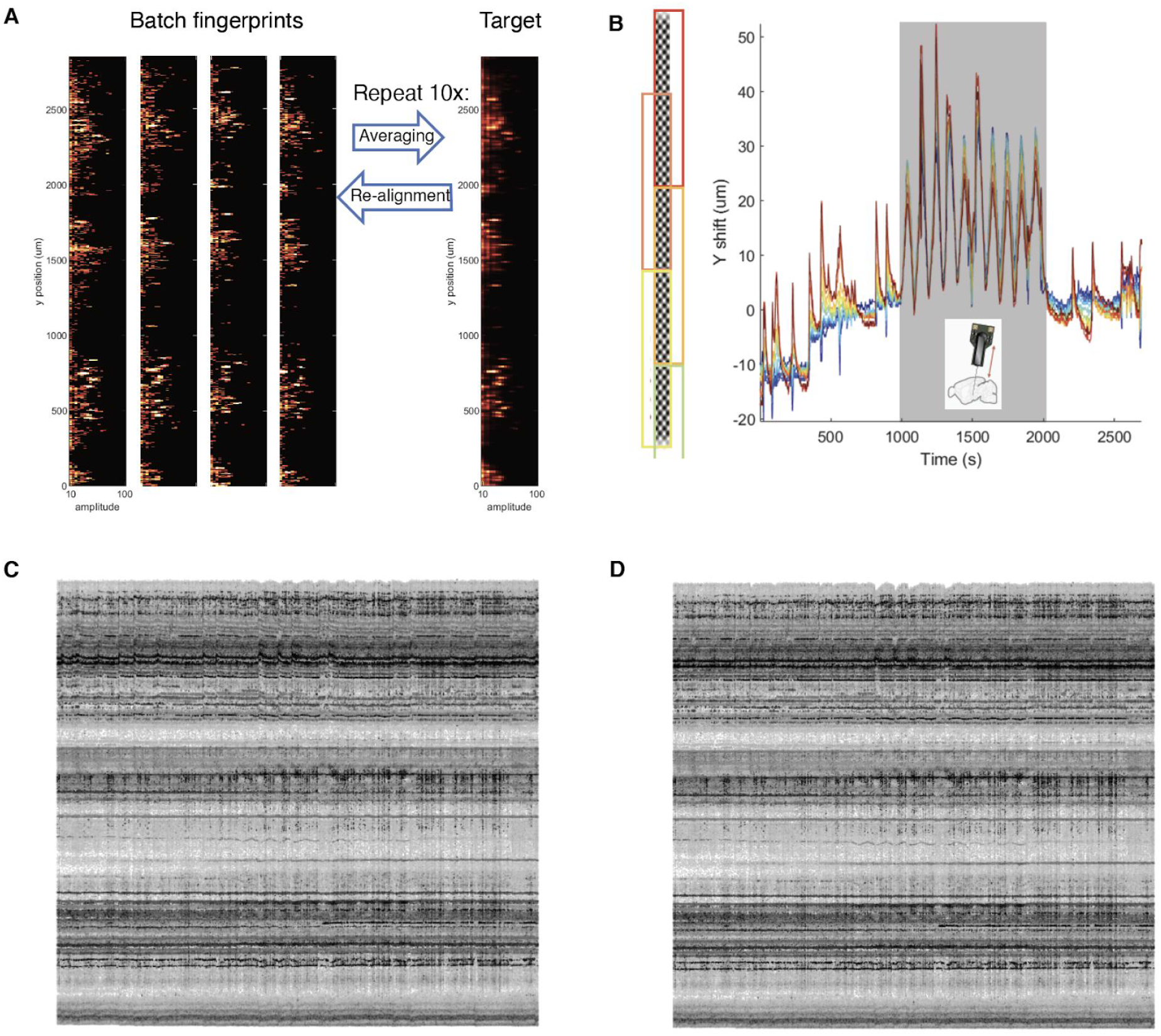
Rigid and non-rigid registration for motion stabilization. (**A**) Four example amplitude/depth histograms from different parts of the recording (like Supp Fig 4C). These provide fingerprints that can be used to align vertically. The average of aligned histograms represents the target image (right), that can be used in another re-alignment pass until convergence. (**B**) (left) Example overlapping blocks used to partition the channels. (right) Estimated drift traces for each block of channels. (**C** and **D**) Drift maps of spikes re-detected from the stabilized binary file. The correction was computed and applied as a rigid registration (**C**), and in a non-rigid manner (**D**). The rigid registration option can be obtained in the software by setting the number of blocks to 1 (see B).

**Supplementary Table 1.** Results of chronic implantation experiments across six labs. Each row represents an individual experimental subject implanted with one or two NP2.0 probes. In the Reason for End column, ‘ED’ means ‘Experimenter’s decision’, i.e. the probe did not fail but the experimenter decided to end the experiment for other reasons. In three cases, ‘ED/Fail’ is indicated because probes exhibited partial failures (described in Notes) but the experiment continued with reduced capability until the experimenter’s decision.

| Lab | Species | Implanted probes | Duration (wks) | Reason for end | Recovery device? | Recovery successful? | Figures | Notes |
| --- | --- | --- | --- | --- | --- | --- | --- | --- |
| Haesler | mouse | One 4-shank | 2 | ED | Y | 1/1 | 2 c,d,e,f, Supp. 1 d |  |
| Haesler | mouse | One 4-shank | 2 | ED | Y | 1/1 | 2 c,d,e,f, Supp. 1 d |  |
| Haesler | mouse | One 4-shank | 2 | ED | Y | 1/1 | 2 c,d,e,f, Supp. 1 d |  |
| Haesler | mouse | One 4-shank | 2 | ED | Y | 1/1 | 2 c,d,e,f, Supp. 1 d |  |
| Haesler | mouse | One 4-shank | 2 | ED | Y | 1/1 | 2 c,d,e,f, Supp. 1 d |  |
| Haesler | mouse | One 4-shank | 2 | ED | Y | 1/1 | 2 c,d,e,f, Supp. 1 d |  |
| Haesler | mouse | Two 4-shank | <1 | ED | Y | 2/2 | 1 e |  |
| Haesler | mouse | Two 1-shank | >8 | ED/Fail | Y | 1/2 | 2 c,d,e,f | One of two probes broke after 8 months |
| O'Keefe | rat | Two 4-shank | >8 | ED | - |  | 2 c,d,e,f |  |
| O'Keefe | rat | Two 4-shank | >8 | ED | - |  | 2 c,d,e,f |  |
| Lee | rat | One 4-shank | >8 | ED | - |  | 2 c,d,e,f |  |
| Lee | rat | One 4-shank | >8 | ED | - |  | 2 c,d,e,f |  |
| Moser | rat | One 4-shank | >8 | ED | - |  | 2 c,d,e,f |  |
| Moser | rat | One 4-shank | >8 | ED | - |  | 2 c,d,e,f |  |
| Moser | rat | Two 4-shank | >7 | ED | - |  | 2 c,d,e,f |  |
| Moser | rat | Two 4-shank | >8 | ED/Fail | - |  | 2 c,d,e,f | One of two probes broke after <1 week |
| Carandini | mouse | One 4-shank | 20 | ED | - |  | 2 c,d,e,f, Supp. 3 |  |
| Carandini | mouse | One 1-shank | 44 | ED | - |  | 2 b,c,d,e,f, Supp. 3 |  |
| Carandini | mouse | One 4-shank | 35 | ED | - |  | 2 c,d,e,f, Supp. 3 |  |
| Carandini | mouse | One 4-shank | <1 | Fail | - |  |  | Probe broke |
| Hantman | mouse | Two 4-shank | >8 | ED/Fail | - |  | 2 c,d,e,f | One shank of one probe malfunctioned |

## Methods

### Neuropixels 2.0 device design

The Neuropixels 2.0 probe consists of one or four shanks (i.e., the thin segment inserted into the brain) and a base (containing the electronics for filtering, amplification, multiplexing, digitization, and power management), fabricated with 130 nm CMOS process as one piece (Wang et al., 2019). The base is affixed to a rigid printed circuit board (PCB) and a thin flexible ribbon cable (“flex cable”) that plugs into a headstage. From the headstage, a 5 m cable runs to a custom PXIe (“Peripheral Component Interconnect (PCI) extension for Instrumentation”; a standardized modular electronic instrumentation platform) data acquisition card (Putzeys et al., 2019) which connects to a computer via an off-the-shelf PXI chassis (e.g. NI 1071, National Instruments), and custom software collects the data and writes to disk. Each of these system components are described in turn below. The details presented here apply to the “alpha” version of the probe, and all data presented in this paper are from this version. A forthcoming “beta” version is planned with broadly similar specifications, but notably with an improved ADC design expected to reduce noise levels. Both the probe electronics (Wang et al., 2019) and the data acquisition system (Putzeys et al., 2019) have been described previously, but aspects of those reports are summarized here for clarity.

The shank is 10 mm long with two columns of sites with 32 μm center-to-center spacing between the two columns, and 15 μm center-to-center spacing along the length of the shank, for 1280 total sites on one shank (Fig 1a), or 5120 sites on the four shank probe version. The shank has a 70 x 24 μm cross-sectional profile. A stress compensation process ensures a deflection of <100 μm from base to tip on each shank. The porous TiN recording sites are 12 x 12 μm^2^ and have an impedance of 148 ± 8 kΩ at 1kHz. The tapered tip of the probe is 175 μm long (tip angle ~ 20°). Due to the planar nature of CMOS manufacturing technologies, the tip is only tapered in the two-dimensional plane of the probe; the tip shape that enters the brain is therefore a line as long as the probe thickness (24 μm). However, the tip can be sharpened to form a point, using a micropipette grinder or similar at an angle as small as 15° (detailed procedure not described here). The triangular tip area is covered by a single large electrode site that can be configured as an internal reference. In addition to the internal reference electrode at the tip, four of the 12 x 12 μm^2^ sites along the shank are also reserved for optional use as internal references, but their use is not recommended because of the much larger impedance of these small sites does not allow for correct cancelation of common-mode noise in all the channels. The center-to-center spacing between shanks on the four-shank version is 250 μm.

The probe base is mounted onto a rigid PCB that has an arrow shape (Fig 1a) with a total width of 3.5 mm at the thin part of the arrow near the shanks, a width of 6.9 mm at the thicker portion, and thickness of ~1.2 mm. The length of the rigid PCB, from shank to flexible ribbon, is ~ 14 mm. The base, affixed to the rigid PCB with wire bonding and epoxy, is 2.2 x 8.7 mm^2^, consumes 36.5 mW of power, and records 384 full band (0.5 Hz - 10 kHz) signals sampled at 30 kHz and at 14 bit resolution. The total data rate is 161.3 Mb/s (or ~23.0 MB/s on disk). The mean input-referred noise level in the action potential range (300 Hz - 10 kHz) is 8.2 μV root mean square, including the electrode noise (the number given in the main text, 7.2 μV r.m.s., reflects the noise of the recording channel alone, which is the only aspect in which the noise of NP1.0 and NP2.0 differ). The input range is 12.5 mV peak-to-peak and the mean gain is ~84. Because of the good channel-to-channel gain uniformity (Wang et al., 2019), gain values are not calibrated for each channel separately, but per probe to reduce global process variation, and this calibration is applied during acquisition. Cross-talk is 0.35% on average between sites at 1 kHz on the single-shank version and 1.51% on the four-shank version. The shank heats < 1° C in the brain.

The flex cable is 43.5 mm long, 4.0 mm wide, and 80 μm thick. Solder pads for attaching referencing and ground connections are provided both near the top of the rigid PCB (Fig 1a, gold squares with holes on the right side of the rigid PCB) as well as along two flexible “wings” on either side of the flex cable; these wings can be cut off if not used. The probe in total (shanks, base, rigid PCB, and flex cable) weighs 0.19 g.

The headstage connects to the flex cable via a 27-pin zero insertion force (ZIF) connector. The headstage has two such connectors, one on either side of the PCB, so that two probes can be connected to a single headstage and stream data from each of their 384 channels simultaneously, for a total of 768 channels. The headstage is 10 x 14.3 mm^2^ in size and weighs 0.72 g. The headstage features a solder pad for ground, and a separate solder pad that connects directly to the tip site(s) and can be used to deliver current. The headstage has a 4-pin Omnetics connector to connect to the cable.

The cable, PXI base-station card, and software are identical to that used for NP 1.0 (Putzeys et al., 2019), but described briefly here for completeness. The cable has two twisted-pair strands (each with 0.41 mm diameter), is 5 m long, weighs 5 g, and terminates in a USB-C connector. Data from all 768 simultaneously recorded channels at 30 kHz are transmitted across this twisted pair cable. The base-station card has four USB-C connectors, accepting input from four headstages (up to 8 probes) simultaneously, and at least two base-stations can be used together in a PXI chassis to stream data to a single computer. A single set of firmware on the base-station allows for recording interchangeably (and simultaneously) from NP 1.0 and 2.0 probes. The base station also accepts an optional digital TTL input channel for synchronization or triggering and an optional battery power supply for isolation. The probe can be configured, and data can be visualized and streamed to disk, with either SpikeGLX (https://billkarsh.github.io/SpikeGLX/) or OpenEphys (https://open-ephys.org/gui) open-source software packages.

As there are 1280 or 5120 electrode sites (i.e. physical TiN electrodes located on the shank) on the single- or 4-shank probe versions, but only 384 recording channels (i.e. signal processing pathways including amplification, filtering, digitization, and data transmission) available, analog switches are used to control which subset of sites is recorded at any given time. The switches are set in software and, after setting, induce a transient voltage deflection lasting < 1 s. The switching schemes that govern which sites connect to which channels differ for the single- and four-shank probe versions, as follows.

The logic of the single-shank scheme is that each block of 32 sites maps onto consecutive groups of 32 channels, allowing for the selection of any contiguous stretch of 384 sites, given that the selected sites start with a site that is a multiple of 32. However, within groups of 32 sites, the mapping from site to channel is scrambled such that any physically clustered group of sites maps onto a set of channels that do *not* correspond to any other physically clustered group of sites (Fig 5a, b). In this way, a given recorded neuron on some set of channels can be localized to one of the groups of sites that connect to those channels according to which of those groups of sites are physically clustered (see also analysis methods section “Combined bank recordings” below and Fig 5).

The logic of the four-shank scheme is that the following selections are possible: any continuous set of 384 channels on any shank; any set that includes 96 continuous sites on each of the four shanks, where the 96 sites are located at the same depth along the probe; any set that includes 96 continuous sites on each of the four shanks, but where the sites on each shank are offset by 96 from shank to shank, forming a diagonal stripe across the four shanks. Other selections consistent with the wiring constraints are also possible (Wang et al., 2019).

### Recording Methods

#### Chronic recordings in mice in Haesler Lab at NERF (Figures 1, 2, Supp Figure 2)

To facilitate chronic recordings with Neuropixels 2.0 probes, we used custom-designed 3D printed fixtures which were developed at NERF, Leuven, Belgium (Figs 1, 2 and Supp Fig 1). Specifically, we used two different types of fixtures, holding either one or two probes respectively. The fixtures consist of two main parts: a) a probe mount and b) a skull connector. The probe mount holds the probe and is reversibly mounted on the skull connector, which is in turn fixed on the animal’s skull. The system further features a headstage cover, which protects the headstage during experiments and a probe retriever tool, which is used to safely recover probes for re-use after an experiment is terminated. All pieces were 3D printed by selective laser sintering (SLS, Materialise, PA12) allowing threaded inserts to be integrated using heat from a soldering iron. For fixture assembly we first aligned the probe(s) in the probe mount with a camera-controlled system and then fixed them in place with light-curable dental cement (SDI Wave). The probe mount holding the probe was then attached to the skull connector with screws. Reference and ground pins on the probe(s) were connected by soldering thin wires both for single and dual probe configurations. Then, the headstage was connected to the probe(s), placed inside the probe mount and covered with the headstage cover. To avoid failures related to connecting/reconnecting the zero-insertion-force (ZIF) connector on the headstage during each experimental session, the headstage was left on the animal for the entire duration of the experiment. The total weight of the assembled fixture with probes was 2.76 g for the dual probe version.

The implantation of the probes was carried out according to animal experimentation protocols approved by the KU Leuven animal ethics committee in accordance with the European Council Directive, 2010/63/EU. Eight male C57BL/6 mice were used for chronic recordings, weighing 28.5 ± 2.4 g (mean + std) at the time of the surgery while 12:12h light:dark schedule was maintained. Two dual probe fixtures with one shank and four shank configurations were chronically implanted into the hippocampus (ML 1.5 mm, AP −2.7 mm w.r.t. bregma) and lateral habenula (ML 1 mm, AP −1.3 mm) equidistant from the midline. Two single probe fixtures with four shank configurations were chronically implanted into olfactory cortices (ML 0.7 mm, AP 2.7 mm).

The animal was anesthetized with ketamine (7.5 mg) and medetomidine (0.1 mg) solution (100 μl/10g body weight). Eyes were moistened with Duratears and hair over the animal’s head was trimmed. Animal’s head was aligned in the anterior-posterior and lateral-medial plane and fixed in the stereotaxic frame by using earbars. Skin was disinfected with iso-betadine Dermicum and ethanol (70%) and ~50 μl xylocaine solution (5% in saline) was injected under the skin over the head for local analgesia. Skin was incised along the midline and removed to expose the skull. Skull was cleaned from remaining biological tissue using sterilized cotton swabs. Skull surface was roughened by scraping grids using a scalpel blade. Vetbond tissue adhesive was applied to cover the exposed tissue. Craniotomy for the desired location was marked with the tip of the drill bit. Head plate was placed over the lambda and secured with Vetbond. Bone screw with a ground pin was implanted over the cerebellum ensuring electrical ground. Headplate and almost the entire skull surface was covered with Metabond leaving the craniotomy location blank. Craniotomy was performed while making concentric circles around the center of the craniotomy and edges were thinned out. Sterile saline solution was applied frequently during the craniotomy to avoid complications. Dura matter was removed by using sharp surgical forceps. Skull clamps were removed and the animal was restrained by the headplate. Probe shank was dipped into a fluorescent dye solution (DiI in ethanol). A wall of 1:1 bone wax and mineral oil was formed around the probe to create a protective medium during the application of cement. After positioning the fully assembled fixture above the target location, it was lowered with the z-axis of the stereotaxic frame with the help of a motorized micromanipulator. The skull connector was fixed in place with Metabond. Then the ground socket on the fixture was connected to the ground pin. A final layer of light-curable cement was applied to cover the remaining gaps. Analgesics (50 μl Metacam, 2 mg/ml) and antibiotics were administered subcutaneously (20 μl Cefazoline, 15 mg/kg body weight) after surgery.

After data collection was completed, we used the retriever tool to recover the probe(s). The retriever tool was first connected to the skull connector. By turning two guiding screws on the retrieving tool, which exert a vertical force on the skull connector, we detached the probe mount from the skull connector. Seven out of eight mounts were retracted successfully without damaging the probes. The probes remained in the probe mount and were cleaned as described previously (Jun et al., 2017). Two out of four probes were re-implanted four and two times, respectively (Supp Fig 2).

#### Chronic recordings in mice in Carandini/Harris Lab at University College London (Figures 2, 4, 5; Supp Figure 3)

Experimental procedures at UCL were conducted according to the UK Animals Scientific Procedures Act (1986) and under personal and project licenses released by the Home Office following appropriate ethics review. The basic implantation strategy has been described previously (Okun et al., 2016). In brief, a mouse with a previously implanted head-plate was anaesthetized and a craniectomy over the left primary visual cortex was made for the probe, and Ag reference wire was cemented in an additional small craniectomy above the olfactory bulb. The probe was then held by a miniature dovetail cemented onto it, and advanced into the brain while recording. When in place, dental cement (Super-Bond C&B; Sun Medical, Japan) was applied to encase the probe PCB and reliably attach it to the skull. Once the cement was fully cured, the dovetail holder was retracted. Next, a custom-designed 3D-printed shell was cemented to the skull next to the probe. This shell served to protect the probe and provided a place to tuck the flex ribbon of the probe and the Ag reference wire into while the mouse was in the home cage. The shell was covered with Micropore tape. The total implant weight was ~3g, and its height was ~2cm. The animal was gradually acclimated to head restraint, starting with a few minutes per day, and monitored continuously for signs of stress. For subsequent recordings, the mouse was head-fixed in front of three screens, and the visual stimuli were shown on the front screen and right screen, while the data was collected. For 4-shank probes, the 384 recording channels were mapped to 96 sites on each shank, forming a horizontal stripe of sites across the 4 shanks. The recordings were made in external reference mode, using the Ag wire or the headplate as the reference signal.

#### Chronic recordings in mice in Hantman Lab at Janelia Research Campus (Figure 2)

All procedures were approved by the Institutional Animal Care and Use Committee at Janelia Research Campus (protocol 19-177). One mouse (male, 13 weeks, Pcp2-Cre X Ai32) was chronically implanted with two four-shank Neuropixels 2.0 probes, one in motor cortex and the other in the cerebellum. The animal was anesthetized with 2% isoflurane and placed on a heating pad in a stereotactic frame (Kopf Instruments). Buprenorpine (0.1 mg/kg) was administered subcutaneously. The head was shaved and sterilized with three alternating swabs of antiseptic scrub (Betadine) and alcohol. The scalp was removed, and the skull was cleaned with a scalpel. A head post (RIVETS (Osborne and Dudman, 2014)) was attached to the skull with UV-curing dental cement (Relyx Unicem, 3M) to provide a flat surface for the implant enclosure and to allow head fixation during recording. A small hole was drilled over right visual cortex, and a stainless steel reference wire soldered to a gold pin was inserted to a depth of 0.5 mm. A drop of silicone sealant (Kwik-Sil, WPI) was placed in the hole, and the gold pin was attached to the skull with UV-curing dental cement (Relyx Unicem, 3M). Craniotomies approximately 1.5 mm in diameter were performed with a dental drill over motor cortex (anterior +0.5 mm, left 1.7 mm w.r.t. bregma) and the intermediate cerebellum (posterior 7.0 mm, right 2.5 mm w.r.t. bregma). The dura was left intact. At each craniotomy, a NP 2.0 probe was slowly lowered to the target depth (2.5 mm into motor cortex and striatum, and 2.5 mm into the cerebellum) with a micromanipulator (Scientifica). The probe was held on a rod with a dovetailed plate at the end, and the probe and plate were kept snug with a set screw. Silicone elastomer (Kwik-Sil, WPI) was applied to the brain surface to seal the craniotomy and to the probe shanks. UV-curing dental cement (Relyx Unicem, 3M) was applied from the top of the probe to the skull to hold the probe assembly in place. Once each probe was secured, the set screw holding it to the micromanipulator plate was loosened, and the fixture was removed. The ground connections for the two probes were patched together and kept in the air. The reference connections for the two probes were patched together and connected to the reference pin in visual cortex. The top of a centrifuge tube (Falcon 15mm, Fisher) was attached to the top of the head post with dental cement to protect the implant. Ketoprofen (5 mg/kg) was then administered subcutaneously, and the animal recovered from anesthesia. The animal received an additional dose of ketoprofen (5 mg/kg) on each of the following two days, and its health was monitored by the experimenter and animal care staff. The animal was gradually acclimated to head restraint, starting with a few minutes per day, and monitored continuously for signs of stress. On each day of recording, data were obtained from the 384 deepest channels on each shank. One of the shanks in the cerebellum failed, so we recorded from four shanks in motor cortex and striatum, and three shanks in the cerebellum. During the recording sessions, the animal was always awake and was either sitting quietly without restraint on the bench or head-fixed.

#### Chronic recordings in rats in O’Keefe Lab at University College London (Figure 2)

These experimental procedures were conducted at UCL according to the UK Animals Scientific Procedures Act (1986) and under personal and project licenses granted by the Home Office following appropriate ethics review. Three male Lister Hooded rats were used for chronic recordings, weighing ~400g at time of surgery and maintained on a standard food deprivation schedule and a 12:12h light:dark schedule. Each rat was implanted with two NP 2.0 probes into the hippocampus and entorhinal cortex. A custom 3D printed metal holder was designed to house the probe and provide a fixation point for the stereotactic surgery frame. No microdrive was used. To protect the electronic components the probe body was encapsulated in epoxy (Araldite Rapid). During the implantation procedure, the probes were secured in the stereotaxic frame, yaw and pitch axes were adjusted to assure that the probe shanks were perpendicular to the horizontal plane through bregma and lambda. A hole for the ground screw was drilled in the right frontal plate and five additional screws were distributed around the implant sites (mEC: 4.3 mm lateral to the midline, 0.3 mm anterior to the sinus; HPC: 2.5 mm lateral to the midline, 4 mm posterior to bregma) to provide anchoring for dental acrylic. The whole surface of the skull except the implant locations was covered with Super-Bond cement (Sun Medical). Two 2.4 mm craniotomies were drilled over the implant locations and bone was carefully removed. The dura was gently lifted and a small incision was made to facilitate the probe insertion. The probes were slowly lowered (10–20 μm/s) until they reached the target location. The brain was sealed with artificial dura (3-4680, Dow Corning). The probe shanks above the brain surface were covered with Vaseline. The probes were then fixed to the skull using dental acrylic. A copper mesh cage was attached to the acrylic and connected to the skull screw to shield the probe assembly from external noise. The probe’s ground and reference were connected to the skull screw. Recordings were made in external reference mode. After experiments were concluded, probes were extracted, detached from the metal holder, recovered, cleaned and reused.

#### Chronic recordings in rats in Moser Lab at NTNU (Figure 2)

All procedures were performed in accordance with the Norwegian Animal Welfare Act and the European Convention for the Protection of Vertebrate Animals used for Experimental and Other Scientific Purposes. Four male Long-Evans rats, aged 3-4 months were chronically implanted with either one or two NP 2.0 4-shank probes. The basic implantation procedure has been previously described (Gardner et al., 2019). Probes were attached to the stereotactic manipulator by means of a custom-designed interface block made from CNC-machined aluminium, to which the probe was glued prior to surgery. Three probes were targeted at left posterior parietal cortex, hippocampus and thalamus (AP −4.0 from bregma; ML 2.8; shanks aligned with coronal plane); the other three probes were targeted at left medial entorhinal cortex (0.1 mm anterior to the transverse sinus; ML 4.3 mm; 25-degree angle in sagittal plane). In two animals, both locations were implanted; in the other two animals, only one location was implanted. All probes were lowered to a depth of 6 mm beneath the brain surface. The probe’s ground and reference terminals were both connected with silver wire to a single ground screw placed over the cerebellum. OptiBond All-In-One adhesive (Kerr) was applied to the skull to provide an adhesive interface layer, and Venus Diamond Flow (Kulzer) and Meliodent dental cement (Kulzer) were used to fix the probe to the skull. The unimplanted portion of the probe shank was enclosed with a protective layer of Vaseline. A plastic screw-top casing made from a 50-ml centrifuge tube was placed over the implant to provide a convenient protective housing.

Electrophysiological recordings were performed while the animals foraged for scattered food crumbs in an open field arena, for a time of 45 minutes or more. Animals were trained to high performance on the task prior to implantation. For each animal, two different channel configurations were recorded: (1) 384 sites on a single shank or (2) 96 sites on each of the four shanks in a horizontal strip. External reference mode was used for all recordings.

#### Chronic recordings in rats in Lee Lab at Janelia Research Campus (Figure 2)

These procedures were conducted in accordance with the Janelia Research Campus Institutional Animal Care and Use Committee. Two male Long-Evans rats, aged ~5 months were each chronically implanted with a NP 2.0 4-shank probe. The implant surgery followed previously described methods (Jun et al., 2017). Briefly, animals were anesthetized with isoflurane and mounted in a stereotaxic frame (Kopf Instruments). After thorough cleaning of the skull, a ground screw was placed through the skull above the cerebellum. A small craniotomy was made above the target areas (AP: 3.24 mm, ML: 0.6 mm in one animal, AP: −5.4 mm, ML: 0.8 mm in the other animal) in the right hemisphere. Probes were centered above the craniotomy and lowered perpendicular to the skull surface to 6 mm depth (rat 1) or at an angle of 20 degrees in medio-lateral direction to 8.1 mm depth (rat 2). Probes were lowered over the course of about an hour to the target depths. Ground and reference of the probe were connected to the ground screw. The craniotomy was covered with artificial dura (Dow Corning Silicone gel 3-4680) and any parts of the probe outside of the brain were covered with sterile Vaseline. The probe was permanently fixed to the skull with dental acrylic, a protective cone made of copper mesh and dental acrylic and light cured cement was built around the probe, and the mesh was connected to the ground. Freely-moving recordings were conducted in a small cardboard box (~ 1 m x 50 cm). Recordings were made in external reference mode.

#### Acute recordings in mice in Dudman Lab at Janelia Research Campus (Figure 1)

Experimental procedures were conducted in accordance with the Janelia Research Campus Institutional Animal Care and Use Committee. A brief (<2 h) surgery was first performed to implant a 3D-printed headplate (Osborne and Dudman, 2014). Following recovery from surgery, the water consumption of the mouse was restricted to 1.2 ml per day. The water restricted mouse was habituated over three sessions to head-fixation and the lick port in the recording setup. The mouse was trained to perform a forelimb reach-to-pull task, similar to the previously described joystick-based reaching task (Panigrahi et al., 2015), for approximately one month. Briefly, the mouse reached and pulled a manipulandum positioned approximately 1.5 cm away from the initial hand position. For acute awake recordings in the fully trained mouse, a small craniotomy was made at 0.5 mm anterior and 1.7 mm lateral relative to bregma in the left hemisphere. The probe was centered above the craniotomy and lowered with a 10 degree angle from the axis perpendicular to the skull surface at a speed of 0.2 mm/min. The tip of the probe was located at 3 mm ventral from the pial surface. The 384 recording sites were uniformly distributed across the four shanks to allow for recording from the approximately 750 μm horizontal and 720 μm vertical tissue area in the dorsal striatum.

#### Acute recordings in mice with optogenetic stimulation in Hausser Lab at University College London (Supp Figure 1)

All animal procedures were performed according to the Animals (Scientific Procedures) Act 1986 under license from the UK Home Office and were approved by the UCL Animal Welfare and Ethical Review Body. Transgenic mice expressing Channelrhodopsin-2 (ChR2) in cerebellar Purkinje cells were generated by crossing a Pcp2-Cre line (B6.Cg-Tg(Pcp2-Cre)3555Jdhu/J) to the Cre-dependent ChR2-EYFP reporter line, Ai32 (Madisen et al., 2012).

To restrain mice during in vivo awake recordings, we installed custom-made aluminium headplates with a 5 mm long and 9 mm wide oval inner opening over the cerebellum. Mice received a steroid anti-inflammatory drug at least 2 hours before surgery (Dexamethasone, 0.5 mg/kg), followed by an analgesic NSAID (Meloxicam, 5mg/kg) immediately before surgery. Inhalation anaesthesia was maintained with 1.5 - 2% isoflurane. The headplate was positioned over the lobule simplex of the left cerebellar hemisphere, angled at 26° with respect to the transverse plane, and attached to the skull with dental cement (Super-Bond C&B, Sun-Medical). Post-operative analgesia (Carprieve, 5 mg/kg) was given for 3 days. After recovering from surgery (minimum 5 days), mice then habituated to the head restraint and the experimental set up for 15 to 30 minutes a day for at least 3 days before the experiment.

On the morning of the recording days, 1 mm-diameter craniotomy and durotomy were performed to allow access for NP probes into the lobule simplex (3 mm lateral to the midline, anterior to the interparietal-occipital fissure). Before the craniotomy, a conical nitrile rubber seal (Stock no. 749-581, RS components) was attached to the headplate with dental cement to serve a bath chamber. The exposed brain was then covered with a humid gelatinous hemostatic sponge (Surgispon) and silicone sealant (KwikCast, World Precision Instruments) until the experiment was performed (1-2 h after recovery).

At the beginning of the experiment, mice were head-fixed, the silicone sealant was removed, and physiological saline solution was immediately applied to keep the craniotomy hydrated. We used single-shank NP 2.0 probes, recording from the 384 most distal channels. To allow post-hoc tracking of the recording site, the electrode shank was coated with a lipophilic fluorescent dye (DiI, Invitrogen). The probe was attached with dental cement to an aluminium dovetail adapter screwed to a 15cm aluminium rod and angled forward 26° (to match the headplate angle and be parallel to the stereotaxic vertical) to target the lobule simplex and lowered at approximately 4 μm/s until reaching the target depth, at which point data acquisition was initiated after 15 minutes of tissue relaxation. For optogenetic stimulation, a fiber-coupled LED generating blue light of 470 nm with a 200 μm diameter and 0.39 NA (M470F3, M95L01 and CFMXB05, Thorlabs) was positioned over the craniotomy. Power at the fiber tip was 4 mW/mm^2^. Optogenetic stimulation consisting of 250 ms steps was used to activate labelled neurons, repeated 50 times at 10 second intervals to minimise desensitization of ChR2. Each recording session consisted of: (1) a 20 minute period of spontaneous activity, (2) a set of 50 optogenetic stimuli, (3) an application of a synaptic blocker cocktail (Gabazine 0.2mM, NBQX 0.8mM, APV 1.6mM) followed by a 20 minute period to let the drugs take effect, and (4) a second set of 50 optogenetic stimuli in the presence of synaptic blockers. After each recording, the probe was soaked in Tergazyme overnight, then washed in distilled water for 1 h and isopropyl alcohol for 1 h.

#### Acute recordings in Steinmetz Lab at University of Washington (Figures 1, 3, 5, Supp Figures 4, 5)

For awake, head-fixed mouse recordings, mice were both sexes, between 2 and 8 months of age. In all cases, a brief (<2 h) surgery to implant a steel headplate and 3D-printed plastic recording chamber was first performed. Following recovery, mice were acclimated over two sessions to head-fixation in the recording setup. During head-fixation mice were seated on a plastic apparatus with forepaws on a rotating rubber wheel. Three computer screens were positioned around the mouse at right angles. On or before the day of recording, mice were again briefly anaesthetized with isoflurane while one or more craniotomies were made with a dental drill. After several hours of recovery, mice were head-fixed in the setup. Probes had an Ag wire soldered onto the reference pad and shorted to ground; these reference wires were connected to an Ag/AgCl wire positioned above the skull. The craniotomies as well as the wire were submerged in a bath of ringer lactate solution, enclosed in the 3D printed plastic chamber. Electrodes were then advanced through the saline and through the intact dura, and lowered to the final position at approximately 10 μm/s. Electrodes were allowed to settle for approximately 20 min before starting recording. Recordings were made in external reference mode. Recording locations included visual cortex, hippocampus, and thalamus.

#### Acute recordings in Svoboda Lab at Janelia Research Campus (Figure 5)

All animal procedures adhered to the guidelines set by the Janelia Research Campus Institutional Animal Care and Use Committee. One wildtype C57BL/6 mouse (male, 6 months) underwent a brief stereotaxic surgery (<2 h) during which a stainless steel headbar was implanted and a recording chamber was built using dental cement for acute head-fixed recordings. Following recovery from surgery, the animal was habituated under head restraint in a recording setup for two days with incremental duration (30 minutes to one hour). On the first day of recording, a 1 mm diameter craniotomy was prepared with a dental drill over the right motor cortex (AP: −0.5 mm, ML: 0.5 mm to Bregma) and dura was left intact. After three hours of recovery, the mouse was head-fixed in recording setup and a single-shank NP 2.0 probe was inserted from the anterior side of the animal at an angle of 47 degrees from perpendicular to the skull surface to the target depth. As such, an insertion covered brain regions including cortex, hippocampus, midbrain, pons and medulla. Prior to each insertion, the tip of the electrode was coated with CM-Dil (Invitrogen), a red fixable lipophilic dye, for later histological reconstruction of probe tracks. Probe was slowly lowered at a speed of 10 μm/s to the final position of 6.6 mm using a micromanipulator (uMP-4, Sensapex, Inc). After reaching the desired depth, the probe was allowed to settle for approximately 10 minutes before commencement of recording. The probe had a wire soldered onto the reference pad which was shorted to the ground pad. This wire was then connected to an Ag/AgCl wire positioned above the skull. During recordings, the craniotomy as well as the reference wire were submerged in cortex buffer (NaCl 125 mM, KCl 5 mM, Glucose 10 mM, HEPES 10 mM, CaCl_2_ 2 mM, MgSO_4_ 2 mM, pH 7.4). Between recording days, the craniotomy was protected with removable silicone sealant (Kwik-Cast, World Precision Instrument). Daily recording sessions lasted 1-2 hours, and were repeated for 2 days in a craniotomy with each insertion separated by at least 200 μm at the point of entry. All recordings were made with open-source software SpikeGLX (http://billkarsh.github.io/SpikeGLX/) in external reference mode and the animal was awake sitting quietly in a body restraint tube throughout the recording sessions.

#### Imposed motion recordings at Steinmetz Lab (Figure 3)

All procedures were approved by the Institutional Animal Care and Use Committee at the University of Washington (protocol 4461-01). To make recordings with imposed motion between the brain and the probe, custom software was used to move the probe after insertion. The probes were affixed to steel rods held by Sensapex uMP-4 micromanipulators (Sensapex, Inc.). Two probes were inserted simultaneously, held by two separate manipulators: a NP 1.0 probe in one hemisphere and a NP 2.0 probe in the opposite hemisphere, using location and angles of insertion matched as closely as possible between the two probes for symmetrical, matched recording sites. The hemisphere used for each probe was counterbalanced across sessions. After ~10 minutes of recording, custom software using the Sensapex API via Matlab was executed to move the manipulators 10 steps forward and 10 alternating steps backwards, each step at 1 μm/sec for a duration of 50 sec, such that the period of the triangle wave motion was 100 sec, the total duration of the pattern was 1000 sec, and the amplitude was 50 μm. A synchronization signal was issued at the start of the motion and at the end for later alignment with neural data. The manipulator position was monitored online programmatically with further API calls during the progression of the motion to ensure that motion proceeded as expected, further confirmed by the visible pattern of motion in the detected location of spikes on the probe (Figure 3d). After completion of the pattern of motion, another ~10 minutes of recording was carried out before ending the session. A total of n=3 such recordings were made, each on a separate day, in n=2 mice.

#### Dual-bank recordings in Carandini/Harris Lab, Svoboda Lab, and Steinmetz Lab (Figure 5)

Subjects were prepared and probes inserted as described above (n=1, Carandini/Harris Lab; n=1, Svoboda Lab; n=2, Steinmetz Lab). Single-shank NP 2.0 probes were used, as the multi-shank version of the probe does not have the scrambled channel mapping required to recover neuron locations. After probes were inserted to the desired depth, ~5-7 mm deep at the tip, three recordings were made for ~10 minutes each: a recording from the sites on bank 1 (i.e. sites 1-384); a recording from the sites on bank 2 (sites 385-768); and a recording from all 768 of these same sites at once, by connecting banks 1 and 2 simultaneously to the 384 readout channels. The recordings were made with SpikeGLX software, which permits the double-bank connection pattern via custom configuration files.

### Data Analysis Methods

#### Spike sorting and quantification of chronic unit yield (Figure 2)

Bandpass filtering (300-9000 Hz) and common average referencing across simultaneously sampled channels (“demuxed CAR”) were applied (using custom software; https://billkarsh.github.io/SpikeGLX/#catgt) to the raw data. We detected events (spikes), calculated spike amplitudes, and calculated firing rates with custom Matlab routines (https://github.com/jenniferColonell/Neuropixels_evaluation_tools), as follows. Putative neuronal spikes higher than the noise floor (constant threshold: 80 μV) were first detected; spikes which occur on neighboring sites within a time window of ± 0.2 ms were grouped together to form an “event”. The spike amplitude for an event was defined as the amplitude on the peak site. Total event/firing rate (Fig. 2d) was then calculated as the sum of all detected events from all channels divided by the time of the entire recording session. As a descriptive function, a linear regression line was fit to the change in total firing rate of each probe/animal days since implantation. The rate of change in log total firing rate was extracted from the slope of the descriptive function. Filled dots in Fig. 2e represent significant correlations (p<0.05; p-values are computed for Pearson’s correlation using a Student’s t distribution using Matlab corrcoef function) of the decay observed in the linear regression line.

For quantification of the unit yield (Fig. 2f), filtered raw data was sorted with the Kilosort2 software (Stringer et al., 2019b). The number of well-isolated single units was automatically estimated by Kilosort2 with a criterion on estimated refractory period violations (<10%). The rate of change in log cluster count was extracted from the slope of the descriptive function (Fig. 2g). Filled dots in Fig. 2g represents significant correlations of the decay, as described above.

#### Motion stabilization algorithm (Figures 3, 4)

To stabilize motion within and across recordings, we developed a data processing pipeline that takes advantage of the increased site density and optimized geometry of the NP 2.0 probes. Here we describe this motion stabilization algorithm in more detail. The pipeline starts by detecting spikes across the entire recording with a new template-based algorithm. We then perform an iterative procedure to simultaneously estimate: 1) the distribution of spike amplitudes across the length of the shank, and 2) the vertical offset of each channel as a function of time and position on the shank. Finally, the vertical offsets are used to shift the data back or “register” it, using an approach based on kriging interpolation. A standard spike sorting algorithm (Kilosort2, (Stringer et al., 2019b)) was then used on the registered data, with different settings for acute and chronic recordings. Below we describe each of the steps in more detail. The algorithm is implemented in Matlab as a part of the Kilosort2.5 package, as well as in Python in an upcoming release.

### Spike template detector

To estimate drift from raw data, we characterize the time-dependent distribution of spike amplitudes and firing rates across the probe length. Since spikes have small spatial footprints on the probe, they can be used to estimate drift at high spatial resolution. In contrast, fluctuations in the local field potential have large spatial correlations, and thus cannot be used to determine micron-scale shifts of the probe.

We start by detecting spikes densely from the entire dataset. One could use standard threshold crossings for this step, but that would not take advantage optimally of the waveform shapes, which are distributed over many nearby channels. We thus devised a more sensitive template-based detection method that we refer to as a “template detector”. Like other template matching algorithms recently used in spike sorting (Pillow et al., 2013; Pachitariu et al., 2016), the template detector sweeps templates across time in a convolutional manner, to find matches with the multi-channel voltage traces. Unlike previous spike sorting methods, the templates we use are not designed and optimized to capture individual waveform shapes from the neurons in the dataset. Instead, they are designed to capture a large range of possible spike shapes, as described below.

Formally, consider a collection of spike templates *W*_*x,y*,σ,*k*_ parametrized by the template center in two-dimensional coordinates (*x, y*) as well as by a spatial scale σ and a template type *k*. The shape of this template on a channel at position (*x′, y′*) is given by:

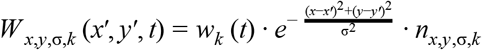

where *w_k_* is one out of a few prototypical template shapes extracted from the data (see below), and *n*_*x,y*,σ,*k*_ is a normalizing constant that ensures ║*W*_*x,y*,σ,*k*_║ = 1. We choose five different σ values of 5, 10, 15, 20, 25 μm to capture spikes of different spatial extent, and we used 2D centers (*x,y*) tiling the probe in the horizontal and vertical dimension, with a horizontal spacing of half the horizontal inter-site distance (16um), and a vertical spacing of half the vertical inter-site distance (7.5um for NP 2.0 and 10um for NP 1.0). Furthermore, we included horizontal centers slightly outside the probe (−16um and 48/64um for NP 2.0/1.0).

To find a good set of prototypes *w_k_*, we detect threshold crossings (six standard deviations below 0) on a subset of the data, ensuring that the crossings are local maxima across channels. We then cluster these one-dimensional waveform shapes using scaled k-means to obtain six representative prototypes *w_k_, k* = 1,…, 6. This ensures that the prototypes are somewhat adapted to the statistics of the dataset, although we expected and noticed that the prototypes are similar across most Neuropixels datasets, probably because sufficiently many distinct waveforms are sampled across a multitude of brain areas.

This procedure gives rise to a total of 57,600 / 69,120 templates for a single Neuropixels 2.0 / 1.0 probe, at a density of 150 / 180 templates per channel. To determine how well a template *W_i_* matches the data *D* at time *t*, we consider the cost function:

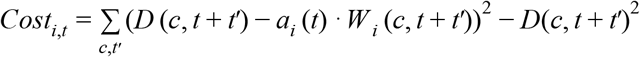

where the sum is over channels *c* and timepoints *t*′ (in our case 61 samples) and *a_i_*(*t*) is the best-fitting scalar amplitude for template *i* at time *t*:

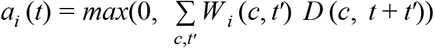

which follows from ║*W_i_*║ = 1 and the additional restriction that all spike amplitudes should be positive. Here, *D* represents the voltage data after high-pass filtering at 300Hz and channel whitening (see (Pachitariu et al., 2016)). It follows that

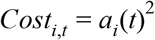

which corresponds to the amount of variance that a template *i* would explain at time *t*. We can thus use 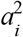 to drive the selection of best matching templates. The local maxima of 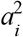 in space and time were then selected as spikes if their amplitudes *a_i_*(*t*) ≥10. The local maximum is over templates that are at most 50 μm away, and at most 20 samples away in time. Examples are shown in Supp Fig 4a.

The calculations to compute *a_i_*(*t*) and its local maxima are performed via custom CUDA code that runs on standard GPUs. Despite using almost 200 templates per channel compared to the more typical 1 template/channel for other spike sorting approaches, the GPU operations are highly efficient due to our chosen factorization of *W_i_* across time and space, running in <10 minutes for a ~1 hour recording on an Nvidia Titan V GPU.

### Estimation of probe drift

The drift stabilization consists of two steps: 1) estimation of drift magnitude as a function of time; 2) interpolation of data according to the drift estimates. In this section we describe the first step, which takes as input the spikes found by the template detector, together with their amplitudes and position on the probe. Since the template positions are restricted to a grid, we further refine the (*x, y*) position on a per-spike basis, by taking the center of mass of the projections *a_i_*(*t*) for all templates *W_i_* = *W*_*x′,y′*,σ,*k*_ with σ and *k* fixed to the values of the identified best-matching template, but *x′, y′* allowed to vary in a ±100 μm range around the values of the best-matching template. Since we only expect drift to vary along the vertical dimension, we ignore the horizontal position of each spike for the rest of this analysis.

Each spike *n* is now described by three numbers: time, vertical position and amplitude or (*t_n_, y_n_, A_n_*). We use these to make the “drift plots” (Figures 3d, 4a and Supp Fig 4b), by assigning darker gray hues to points of higher amplitude (Supp Fig 4b). These plots show clear horizontal bands where spikes of high amplitude are likely coming from a single well-isolated neuron, or a collection of neighboring neurons with similar amplitudes. The goal of drift estimation is to track the position of these bands across time, in a way that is either 1) consistent across the probe (rigid estimation) or 2) varying as a function of depth (nonrigid estimation). We describe a method to determine rigid shifts for a section of the probe, and then describe how the method can be extended to estimate nonrigid shifts across an entire probe.

#### Rigid drift estimation

We start by splitting the dataset into temporal batches that tile the entire recorded duration (Supp Fig 4c). The batches for all analyses in this paper have been set to ~2 sec, but the software allows setting longer or shorter batches, with some caveats that we discuss at the end of this section. For each batch, we convert the data representation from a collection of spikes (*t_n_, y_n_, A_n_*), to a single 2D histogram of depths *y_n_* and amplitudes *A_n_*. We chose equally spaced bins for depth (5 μm), and logarithmically-spaced bins for amplitude (20 bins from the lowest to highest amplitudes, in our case 10 to 100). We then log-transformed the counts 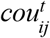 in each bin *i,j* for each batch *t* according to 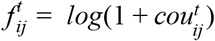, where *i* indexes the depth bins, and *j* indexes the amplitude bins. Finally, we applied a gaussian smoothing filter with a standard deviation of 0.5 bins in each direction to get the final representation 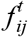 (Supp Fig 4c).

The matrices 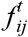 can now be used for vertical registration, because they contain the fingerprint of neural activity patterns across the probe length. The objective of registration is to align these fingerprints to each other vertically so that they match each other best. To achieve this objective, we use a standard correlation-based registration approach, similar to the approach used for 2p calcium imaging data (Foroosh et al., 2002; Pachitariu et al., 2017). The process needs a registration target to align each frame to. One could use a single 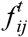 matrix as the target, but that would not take full advantage of the SNR of the data, because individual 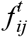 matrices are noisy. Instead, we jointly optimize a registration target *F*, and the registration offsets *d_k_* which are needed to shift each matrix 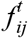 so that it best matches *F*. We iteratively optimize the following cost function:

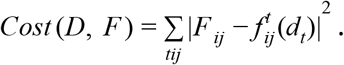

On each iteration, we first hold *F* fixed and optimize *d_t_* individually for each batch, and then hold *d_t_* fixed for all *t* and optimize *F* (Supp Fig 5a). The optimization of the second step simply sets 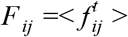 where < > is the averaging operator. For the first step, we calculate the cost function reduction for each integer value of *d_t_* in a reasonable range (± 20 bins). On consecutive iterations, we remember *d_t_* from the previous iteration and refine it in a range around its previous value. This process is initialized with *F* = *f^t^* for an arbitrarily-picked batch *t* which we choose to be in the middle of the recording duration. The optimization needs 5-10 iterations to converge.

At convergence, we obtain the best estimates of *d_t_* to within 5 μm vertical precision, which we further refine using an interpolation-based method that upsamples the curves of correlation between 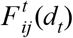 and *F* at different values of *d_t_* (individual curves in Supp Fig 5b). For upsampling, we use kriging interpolation with squared-exponential kernels of standard deviation of 1 bin.

We note that the estimation of drift has a space/time accuracy tradeoff. The signal-to-noise ratio (SNR) of the drift estimator varies strongly with the total number of spikes in the batch, regardless of which channels those spikes were recorded on. Thus, one can either get a temporally-precise estimate of drift using all channels on a Neuropixels probe (384) or a spatially-precise estimate for smaller sections of the probe, but using longer batch durations. For 32 electrodes, the minimum useful batch duration could be on the order of tens of seconds, although a less precise drift estimate can probably still be obtained with shorter batches. Determining the optimal trade-offs between temporal and spatial resolution of drift is a topic of further work. Furthermore, there is likely more information available in the multi-dimensional features of single spikes than the amplitude features we have used here.

In the next section, we assume that drift can be estimated relatively well for subsections of ~1/3 of the probe for a batch duration of 2s, and then introduce a smoothing operation to ensure that even segments with few spikes are assigned good drift estimates.

#### Nonrigid drift estimation

Because one group of 384 sites on a NP2.0 probe is ~3 mm long, it is possible that drift is different in different segments of the probe, requiring nonrigid registration. This type of nonrigid movement can be seen clearly in the original drift plots (Supp Fig. 4b) and also in the drift plots recomputed on rigidly-corrected data, where it appears as residual drift (Supp Fig. 5c). From our experience, it is common for cortical and subcortical regions to have differential drift, although this could depend on the penetration angle and other factors specific to each preparation.

To correct for nonrigid drift, we again turn to a common practice in image analysis: splitting the data into multiple segments and assuming approximately rigid drift for each small segment (Pachitariu et al., 2017). For a complete Neuropixels probe, we use 7 segments that have consecutive overlaps of 50% of their lengths (Supp Fig 5b). In other words, each channel is included in exactly two consecutive segments. For each of these segments, we run the rigid drift estimation algorithm above, with one modification that arises before the upsampling step. This modification is a simple smoothing operation across registration blocks, timepoints and registration offsets, which we use to increase the SNR of the curves of correlation between 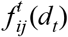 and *F*, now computed separately for each segment. For smoothing we use a Gaussian kernel of standard deviation of a half sample in each direction. We found this smoothing operation to be often necessary, because each block has a relatively small number of channels and timepoints. Note it is preferable to improve SNR *before* taking the peak of the cross-correlation curves. This is because the nonlinear nature of the *argmax* operation can result in outlier drifts for some samples, and smoothing would then be highly detrimental to nearby samples. After smoothing the cross-correlation curves, the estimation process for the best shifts proceeds like in the rigid case, including the upsampling step (Supp Fig 5b).

Finally, we upsample the drift estimates for the 7 segments of channels to produce a private estimate for each channel. This is done by assigning the drift estimate for each segment to the mean channel in that segment, and using a standard interpolation method (Akima spline interpolation). The final result of nonrigid drift estimation is a set of drift estimates parametrized as *d*(*k, t*) for each channel *k* and each batch *t*, that we use in the next phase to stabilize the data.

### Drift correction using kriging interpolation

Given the drift estimates, we must correct the data by shifting it up or down on each batch and each set of channels. We directly modify the high-pass filtered and channel-whitened version of the data, since this is what we will use for spike sorting. Each sample time point *T* with data *q_T_* inherits the drift estimates *d_t_* for the batch *t* it belongs to, where *q_T_, d_t_* are understood to indicate vectors across channels. Each channel in turn is assigned a horizontal and vertical position defined as *x* and *y* respectively. Our goal is to estimate the data at channel positions *x* and *y* + *d_t_*, which we do using kriging interpolation. This is a method for producing interpolation matrices that assumes the data was generated by a Gaussian process, which we choose to have generalized Gaussian kernels with a spatial constant σ defined by the user (15 μm by default). The joint covariance of the sites *z*_1_ = (*x,y*) and *Z*_2_ = (*x, y* + *d_t_*) can be written as:

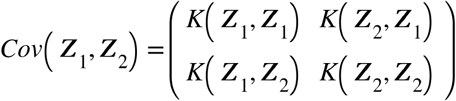

where *K_p_* (*Z*_1_, *Z*_2_) = *e*^−*E*(*Z*_1_*Z*_2_)^*p*^/σ^*p*^^ is the kernel and *E*(*Z*_1_, *Z*_2_) is the matrix of Euclidian distances between all pairs of two-dimensional coordinates in *Z*_1_ and *Z*_2_. Given the observation of data *q_T_* on channels *Z*_1_, we can use the covariance to obtain the predictive distribution on channels *Z*_2_, which has a mean given by:

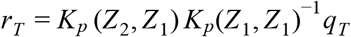

with *r_T_* being the registered data. The kernel computation and matrix inversion need to be calculated once for each data batch, making this interpolation method computationally efficient on GPUs. We also added a small constant to the diagonal of *K* (*Z*_1_, *Z*_1_) to make the inversion operation more stable, and used a default *p* = 1 for all analyses in this paper.

After registration, we rerun the spike detector algorithm to make new drift maps (Supp Fig 5c,d). Some small residual drift can be seen for the rigid registration algorithm (Supp Fig 5c), while the nonrigid algorithm corrects the drift even better (Supp Fig 5d). The spike sorting algorithm should be robust to potential small amounts of residual drift. Note also that one putative neuron in fact appears to move *more* after drift correction than before (compare Supp Fig 5c,d to Supp Fig 4c). We believe this could reflect a neuron stuck to the probe and moving up and down with it, a reminder that drift alone does not account for *all* non-stationarities in a neural recording.

Finally, we took special care of modifying the data at batch boundaries, where an abrupt change in the registration offsets computed by the algorithm would lead to apparent discontinuities in the data, that in turn can affect the spike sorting algorithm. To minimize these discontinuities, we use an overlapping window of 64 samples between consecutive batches, and interpolate linearly between the previous and next batch, with coefficients that vary linearly from 0 on the first sample to 1 on sample 64 of the overlap window. Note that we use a similar approach to minimize discontinuities due to high-pass filtering. The final result of this step is a high-pass filtered, channel-whitened and drift-corrected data file of the same size as the original data and which can be further used for spike sorting with, in principle, any algorithm.

### Spike sorting with Kilosort2

After the data has been registered, we proceed to spike sorting using the Kilosort2 clustering and template matching algorithm. Because the drift has already been corrected, we did not need to use the drift tracking step in Kilosort2. The templates were found by processing the data batches in pseudo-random manner, and then frozen to their final values for the entire template matching step, instead of being allowed to change like in Kilosort2. Although we did not use the tracking step of Kilosort2, we do take advantage of other improvements in Kilosort2, like the initialization of new templates from the residuals of the template matching algorithm, and the post-processing steps that split and merge the final templates to create units of higher quality. The pre-registration step enables the use of powerful quality control (QC) criteria to further automate the spike sorting pipeline. For example, the isolation distance between clusters in feature space would not be a good QC test if the features of the clusters are drifting over time and thus getting smeared in feature space. However, if we assume the data is stationary, the isolation distance should become more widely useful. Post-processing packages already exist that compute this metric, and we plan to incorporate such a metric into a Python-based version of Kilosort (coming soon).

### Chronic recordings

For chronic recordings, we tried to concatenate the raw data files and run the Datashift algorithm as described above. The results were not always satisfactory, especially when the recording sessions were separated by weeks. In these cases, we noticed substantial changes in the drift maps over days that were not consistent with a simple vertical drift but may have instead reflected neurons becoming more or less active over days (Deitch et al., 2020). To analyze these data, we used a different strategy. We started by estimating drifts on each session separately and then we aligned the registration targets from each session to each other, adding this additional drift to the per-session drift estimates. We rounded these drift estimates to integers, and shifted data from both sessions by these amounts to approximately align the recordings. Finally, we ran the standard Kilosort2 algorithm with drift tracking, allowing the waveforms to change as a function of time during template matching.

#### Acute motion stabilization test, data analysis (Figure 3)

To analyze the success of the acute motion stabilization algorithm, we computed three metrics of success: number of stable units, number of unstable units, and correlation coefficient between firing rates and motion trajectory. To compute stable and unstable unit counts, we ran the Kilosort2 algorithm on the datasets before and after motion stabilization. Units were selected for analyses if they passed two stringent criteria: 1) the spatial extent of the waveform was sufficiently small; 2) the fraction of refractory period violations was <10% from that expected of a Poisson process with the same mean firing (Stringer et al., 2019b). For the most part, criterion 1) distinguished neuronal waveforms from noise, and criterion 2) distinguished well-isolated single units from the background activity. We also excluded any units with firing rates less than 0.1 Hz. To compute the correlation coefficient between firing rates and motion trajectory, we binned spikes (1 s bin size) across the period of the recording with imposed motion and computed the correlation coefficient with the triangle wave motion pattern. Since either positive or negative correlations would indicate the presence of problematic failures to detect spikes at some phase of the motion, we took the absolute value of the correlation coefficients. To estimate the null distribution for this measure, we computed a ‘shuffle’ correlation between the firing rate during a period without imposed motion and the triangle wave motion pattern (Fig 3f, ‘shuffle control’). Since firing rates during that period could not truly vary with the imposed motion, this control provides a lower bound on the measure. We used the null distribution to establish 95% confidence intervals, based on which we classified units as stable/unstable.

#### Chronic cross-session stabilization test, data analysis (Figure 4)

To assess the stability of the sorted units in pairs of spliced recordings performed on different days, we used units whose baseline firing rate between the two days differed by no more than 4-fold (thus 25%±16% of the units were discarded, mean ± standard deviation across shanks). Note that true baseline firing rates in mouse visual cortex vary substantially over long timescales (Deitch et al., 2020), so these excluded units may not represent recording instability. Next we tested whether each unit’s visual responses differed significantly across the 112 natural images that were shown (p-value < 0.01 in a Kruskal-Wallis test of the visual responses). Units that satisfied both criteria were considered to have a reliable visual fingerprint response. The fingerprint consisted of the average spike count response to each image and the mean (across all images) response PSTH.

The first day visual fingerprint of a unit was compared to its own second day visual fingerprint and to the second day visual fingerprint of another unit whose location on the probe was closest to the original unit. The comparison was performed by correlating the spike counts and PSTHs of the two fingerprints, then taking the average of the two values. Such comparison can have two possible outcomes: (i) match - when the fingerprint on the first day has higher similarity to the unit’s own fingerprint on the second day; (ii) mismatch - when the fingerprint on the first day has higher similarity to the fingerprint of the other unit on the second day.

The ratio between the number of matches and mismatches of visual fingerprints allows us to infer the number of stably-recorded units, based on an approach we described previously (Okun et al., 2016). Briefly, using the equation for total probability, we have:

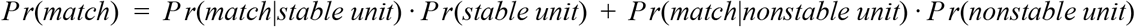

Since *Pr*(*nonstable unit*) = 1 – *Pr*(*stable unit*), we get that

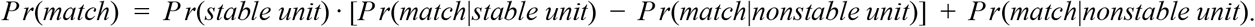

Thus,

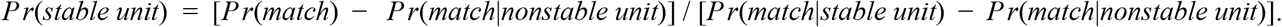

Finally, we use the fact that *Pr*(*match|stable unit*) ≤ 1 and *Pr*(*match|nonstable unit*) ≤ 1/2 (if the neuron corresponding to the unit on the second day is different from the original neuron, match is no more likely than mismatch) to derive that *Pr*(*stable unit*)≥2*Pr*(*match*) – 1. For the same reason, *Pr*(*match|stable unit*) ≈ 1 and *Pr*(*match|nonstable unit*) ≈ 1/2 imply *Pr*(*stable unit*) ≈ 2*Pr*(*match*) – 1.

The above equation (inequality) was used to convert the observed match probability on each shank into an estimate of the (lower bound on the) probability that the units recorded on the shank are stably tracked.

*Pr*(*match*) was taken as the proportion of well-isolated units that matched out of all well-isolated units output by the algorithm for shift-corrected spike sorting (a version of Kilosort 2 (Stringer et al., 2019b)). A unit was considered well-isolated if Kilosort2 labeled it as ‘good’ rather than ‘mua’ (48% of all units with a reliable visual fingerprint response). In a subset of recordings we compared these results to results obtained after manual curation of the output using Phy (Rossant et al., 2016). We found that manual curation produced results that were quantitatively almost identical to those obtained by performing the analysis on the immediate output of the automated spike sorting algorithm (data not shown).

#### Combined bank recordings, data analysis (Figure 5)

We performed spike sorting on the recordings from each bank (1, 2, and 1+2, as described above) using the standard Kilosort 2 algorithm (Stringer et al., 2019b). The template waveform of each unit returned by Kilosort 2 was analyzed to determine whether the template could be included and to determine the mismatch score on each bank. The mismatch score was defined as:

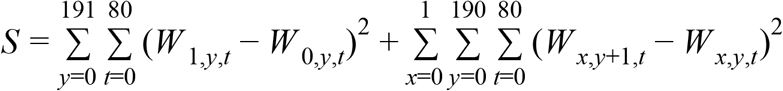

where *W_x,y,t_* is the waveform on the channel at coordinate (x,y) at time t relative to the spike. The coordinates x and y specify the two columns (x) and the 192 rows (y), and are sorted differently for each bank according to the channel mapping. Units were excluded from this analysis when they were not labeled ‘good’ by Kilosort 2, when they had template waveforms extending across more than 20 channels (defined as the number of channels with amplitude at least 10% of the maximum amplitude on any channel; values >20 indicate low amplitude spikes that were difficult to discriminate), and when they had mismatch scores that differed by less than 0.5 between the two banks.

The change in SNR between single- and combined-bank recordings was computed by first calculating the r.m.s. voltage in each case, after band-pass filtering the raw data between 300 and 10000 Hz with a 3-pole Butterworth filter (r.m.s. in each case for an example recording shown in Fig 5e). Then, we assumed that the spike amplitude decreased by a factor of 2 for spikes recorded on any given channel, and computed the resulting change in SNR on each channel using this 2x decrease in signal with the empirically measured changes in ‘noise’ (r.m.s. voltage).

## Acknowledgements

We are grateful for funding from the Wellcome Trust (204915/Z/16/Z), the Klingenstein-Simons Foundation (N.A.S.), the Pew Biomedical Scholars program (N.A.S.), the National Institutes of Health (1U01NS113252-01 to N.A.S.), the Research Foundation Flanders (FWO) (G096219N, to S.H.), KU Leuven (C14/17/109 to S.H.), the Research Council of Norway (National Infrastructure Scheme, NORBRAIN, grant number 295721; FRIPRO grant number 286225; Centre of Excellence grant number 223262 to E.I.M.), the Kavli Foundation (E.I.M.), the BBSRC (BB/P020607/1, to M.O.), the Academy of Medical Sciences and Wellcome Trust (Springboard SBF002\1045 to M.O.), the Sainsbury Wellcome Centre for Neural Circuits and Behaviour (A.L.), and the Hermesfonds with a VLAIO Baekeland mandate (HBC.2018.2114 to R.v.D.).

## Author Contributions

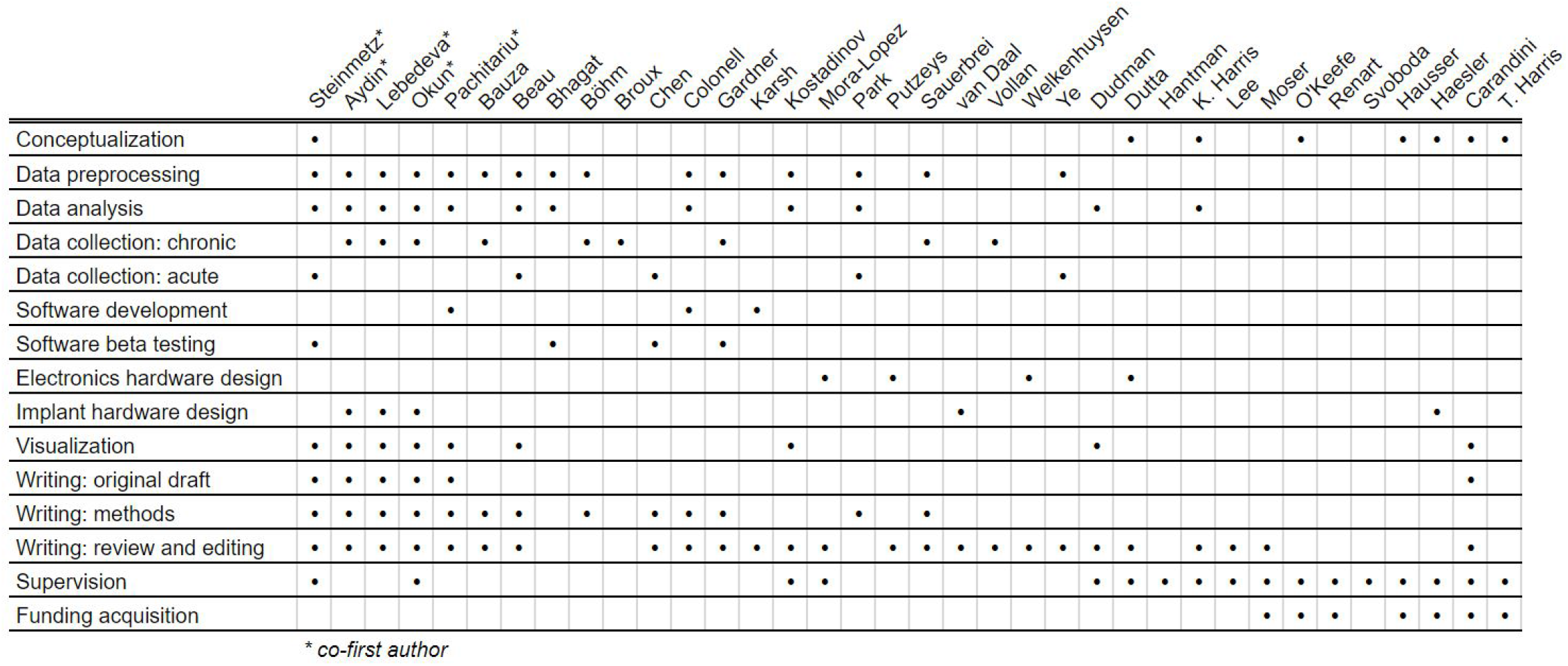

